# Repurposing passenger amplifications as Trojan horses identifies MPZL1 as a potent target for solid cancers

**DOI:** 10.1101/2025.09.17.675062

**Authors:** Sonia Jiménez-Vázquez, Christos Patsis, Mara Mitstorfer, Melanie Grimm, Luise Butthof, Lena Wendler-Link, Lio Böse, Hendrik Wiethoff, Ilse Hofmann, Kai Breuhahn, Thomas Longerich, Matthias M. Gaida, Peter Schirmacher, Judith Feucht, Marco Breinig, Darjus F. Tschaharganeh

## Abstract

Personalized cancer therapies focus mostly on targeting driver alterations, such as oncogenic point mutations or oncogenic driver events within large somatic copy number alterations. However, these alterations are often not actionable or only present in a small subset of patients. We hypothesized that passenger events, specifically in amplified regions, could be therapeutically exploited by providing actionable molecules on the cell surface, serving as Trojan horses for specific therapy delivery. Applying a multiomics *in silico* approach, we identified *MPZL1* (*Myelin protein zero-like 1*), a glycosylated cell surface receptor located on chromosome 1q, as a promising candidate, which is amplified in up to 75% of cases in several solid cancers. Notably, immunohistochemistry of a wide range of human cancer tissues (n=2244 samples) as well as normal tissues revealed strong membranous MPZL1 expression in a majority of solid tumors (e.g. 48% of hepatocellular carcinomas or 89% of triple-negative breast cancers), whereas healthy tissues were mostly negative or just faintly positive for MPZL1. Next, we generated a highly specific monoclonal antibody directed to the extracellular domain of human MPZL1 protein and utilized this antibody to produce MPZL1 CAR-T cells. MPZL1 CAR-T cells showed high specificity as well as high sensitivity in targeting a multitude of human cancer cell lines (e.g. liver, breast, and lung cancer) with high MPZL1 expression *in vitro.* Finally, we demonstrate strong therapeutic efficiency of MPZL1 CAR-T cells not only in different human xenograft tumors *in vivo* but also in a unique autochthonous liver cancer mouse model. Our work provides a framework to target passenger events within large chromosomal amplifications, reveals MPZL1 as a new trojan horse entry point for therapies of 1q-amplified cancers, and as such opens a new avenue for innovative approaches in anti-cancer drug development.

## Introduction

Somatic genetic alterations are the underlying cause of cancer and consequently they are widely used to define respective therapies for precision medicine. Most therapies are focusing on driver genes, which are fueling the progression of tumors and thus are attractive targets for cancer therapy [1], [2]. In particular point mutations in specific oncogenic driver genes, e.g. *KRAS* or *BRAF*, have been therapeutically exploited by developing corresponding inhibitors, which show great promise in patients harboring cancers with these mutations [3], [4], [5], [6]. Moreover, driver genes, which are amplified by somatic copy number alterations (SCNAs), and thus exhibit increased dosage of their genetic material, are also targets of drug development efforts [7]. The most prominent gene in this regard is HER2/ErbB2, which is amplified in approximately 15% of breast cancer patients [8], and can be targeted either by small molecule inhibitors or by monoclonal antibodies and antibody-drug conjugates [9], [10]. However, although we have made great progress in precision cancer therapy, there are still approximately two thirds of cancer patients for which no precise molecular target exists (e.g. due to druggability) and therefore are not amenable for precision oncology [11].

More recently, many efforts were undertaken to circumvent the need to target driver mutations directly. Many of them are using the concept of synthetic lethality (SL), in which dependency on a specific pathway or gene is gained in cancer cells due to their genetic constellation, even independent of a genetic driver event [12]. An example of SL exploring point mutations is provided by the gene pair PARP and BRCA1/2, where inhibition of PARP results deleterious for BRCA-deficient cells, but does not affect BRCA wild-type cells [13], [14]. Moreover, collateral lethality is a subtype of SL which specifically targets genes associated with passenger deletions [15], and often refers to paralog pairs such as ENO1-ENO2 [16] or ME2-ME3 [17]. We are currently exploring this field and have identified a novel SL gene pair, MFRN1-MRFN2, in which targeting MRFN2 induces cell death in chromosome 8p deleted cancers [18]. Although these findings are promising, their implementation into clinical practice is still hindered by the challenging development of specific inhibitors for most of the emerging targets, as well as their testing in clinical trials.

Whereas targeting of bystander alterations in large genomic deletions is already widely accepted, approaches harnessing collateral vulnerabilities created by genomic amplifications have only emerged recently [19], [20]. It is worth noting that large genomic amplifications do not only comprise driver genes but also lead to dosage elevation of many other genes [7], which could potentially be exploited for novel therapeutic strategies. Here, we hypothesized that cell surface proteins located in amplified genomic regions could serve as specific entry points for therapies targeting cancer cells –analogous to Trojan horses– regardless of their biological function. The amplification of these genes and the corresponding increase in protein expression in cancer cells could create a therapeutic window that allows selective targeting of cancer cells while sparing normal cells, thereby contributing to precision medicine.

## Results

### An integrative *in silico* approach identifies MPZL1 as a potential targetable passenger amplification

To compile a comprehensive list of cell surface proteins (CSPs), we employed an open-access resource containing 1492 human membrane-bound proteins [21]. We annotated the genes coding for these CSPs and examined their copy number status in a publicly available dataset of 2703 pan-cancer patients from a joint study by the International Cancer Genome Consortium and the Cancer Genome Atlas Network (ICGC/TCGA) [22]. This analysis revealed that 90 genes coding for CSPs were amplified in up to 30% of the profiled tumors (Figure 1A) (Suppl. Table S1). Interestingly, the most frequently amplified genes were all located on the long arms of chromosomes 1 and 8 (Figure 1B). Since the long arm of chromosome 1 (chr. 1q) harbored the majority of these genes (N=77), we concluded that chr. 1q is a hotspot for amplified cell surface protein-coding genes in cancer, and focused further on this chromosomal region for subsequent analyses.

**Figure 1.**
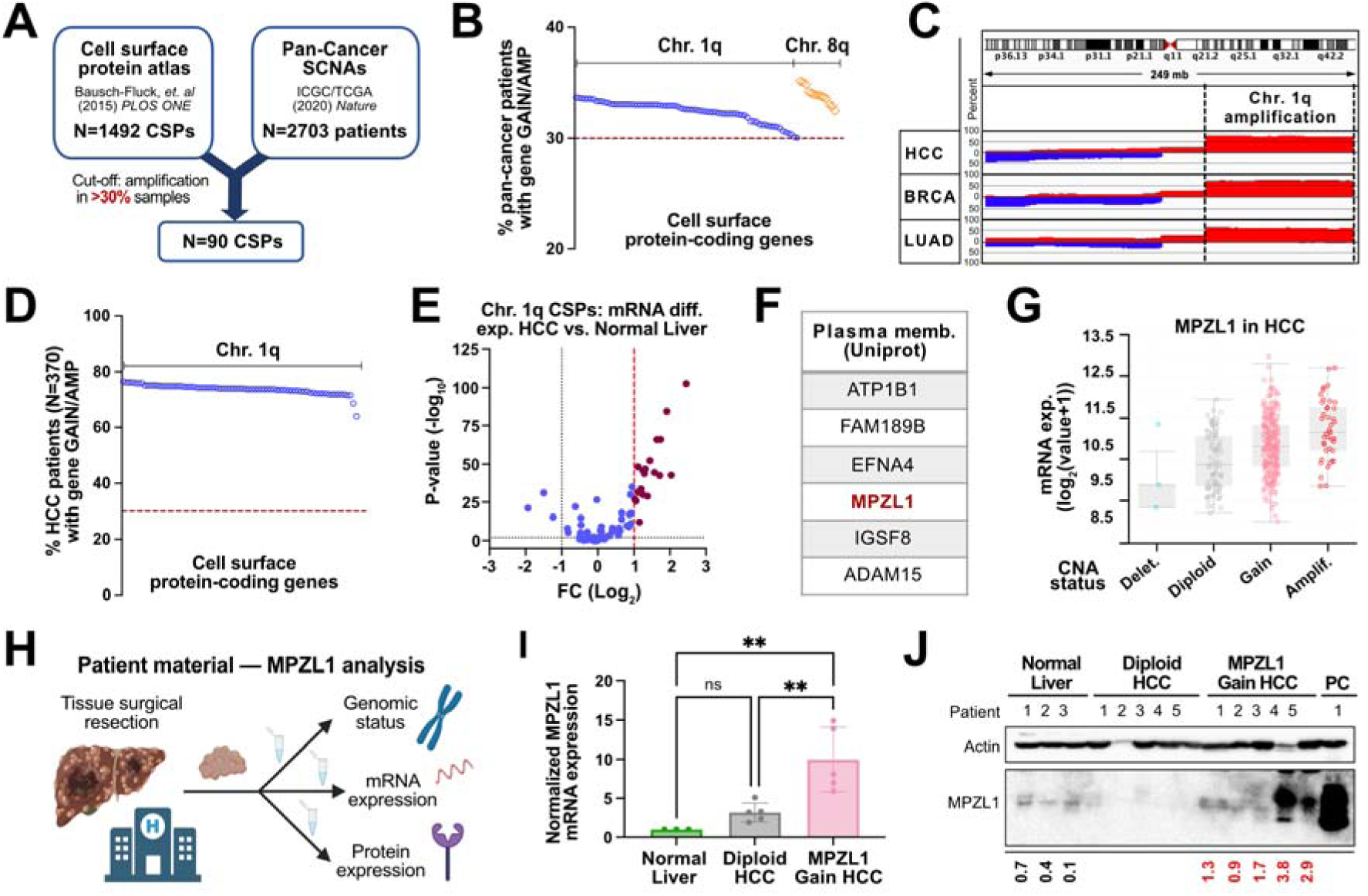
Identification of MPZL1 as a potential targetable passenger amplification. **(A)** Schematic of the *in silico* screen performed. The CNV status of the genes coding for the CSPs compiled in the CSPA (N=1492) is surveyed in a pan-cancer dataset (N=2703 patients), using the online tool cBioPortal. CNV status is categorized by the GISTIC2.0 algorithm, which assigns a specific score to each gene in each patient: -1 (shallow deletion), 0 (diploid), 1 (gain), and 2 (amplification). In this study, hits are defined as genes that receive a score of 1 or 2 in more than 30% of samples (N=90). **(B)** Chromosomal arrangement of hits is analyzed using the *BioMart data mining tool* from Ensembl, with chromosome 1q identified as a hotspot for CSP-coding genes (N=77) across cancers. (**c**) Genome view of recurring patterns of SCNAs across tumor entities from publicly available TCGA datasets (HCC: hepatocellular carcinoma; BRCA: breast invasive carcinoma; LUAD: lung adenocarcinoma). The data is presented as the number of samples in percentage with log_2_(CN/2)>0.1 for amplifications (red) and log_2_(CN/2)<-0.1 for deletions (blue) (HCC: n=370; BRCA: n=1080; LUAD: n=516). **(D) – (J) Downstream analyses are focused on HCC. (D)** Percentage of HCC patients (n=370) who exhibit a gain (score 1) or amplification (score 2), as defined by the GISTIC2.0 algorithm, of each of the hits. Analysis performed with cBioPortal. **(E)** Volcano plot depicts mRNA differential expression data of chr. 1q frequently amplified CSP-coding genes in liver tumors compared to normal liver tissues (Tumor:n=369; Normal:n=160). Analysis done using the GEPIA2 online resource and the ANOVA method. Data is expressed as log_2_(fold-change) with cut-off set at log_2_(FC)>1 (N=19 hits). **(F)** List of CSPs annotated in UniProt as being preferentially found at the plasma membrane. MPZL1 CSP is selected as a potentially targetable bystander amplification. **(G)** Boxplots show the correlation of MPZL1 CNV status –as defined (A)– and its mRNA expression levels in the HCC TCGA dataset (n=360). mRNA data is quantified using the expectation-maximization algorithm and expressed as log_2_(value+1). Analysis performed using cBioPortal. **(H) – (J) MPZL1 in human liver samples from patients. (H)** Schematic of the workflow. Created with *BioRender.com*. **(I)** MPZL1 mRNA expression is evaluated via microarray analysis. **(J)** MPZL1 protein levels are assessed via immunoblotting, with actin used as loading control. Numbers below the blot represent quantification of protein bands, where the intensity of MPZL1 is normalized to the intensity of actin via densitometry-based analysis using ImageJ.

To ascertain the prevalence of chr. 1q amplification in different solid tumor types, we analyzed copy number data of publicly accessible datasets across different cancer entities within the TCGA Firehose Legacy study [23]. We found that more than 70% of tumors in patients with hepatocellular carcinoma (HCC) [24], invasive breast carcinoma (BRCA) [25], and lung adenocarcinoma (LUAD) [26] exhibit such amplification (Figure 1C), which spans approximately 100 mb, extending from segments 1q21.1 to 1q44. Consistently, the identified CSPs located on this region are amplified in over 70% of samples across the HCC, BRCA and LUAD cohorts (Figure 1D, Suppl. Figures S1A and D) (Suppl. Table S2). Thus, due to the frequency and omnipresence of this amplification, a therapy directed to a CSPs on chr. 1q would broaden its impact to several solid cancers.

To further define promising targets we profiled expression levels of CSPs in healthy tissues and their malignant counterparts to minimize off-target toxicities. We first focused on HCC, as this tumor type exhibits the highest amplification levels chr. 1q, and performed differential mRNA expression analysis of tumor tissue relative to healthy tissues using the web-based tool GEPIA2 [27]. 19 chr. 1q-located CSP-coding genes were found to be at least 2-fold higher expressed (|Log_2_FC|>1) in liver tumor tissues compared to healthy tissues (Figure 1E) (Suppl. Table S3). Remarkably, conducting differential mRNA expression analysis in breast and lung tissues mirrored the results obtained in HCC (Suppl. Figures S1B and E). Additionally, an ideal therapeutic target should exhibit high expression levels at the plasma membrane. To ensure this, we surveyed public information available about the 19 CSPs in different protein databases [28], [29] and observed that only six of the candidates were described to preferentially locate to the plasma membrane (Figure 1F) (Suppl. Table S4), while the remaining were reported to locate to other membrane compartments, primarily associated to the endosomal secretory pathway, such as the Golgi apparatus, lysosomes or endosomes. Interestingly, while all these CSPs have been implicated in cancer processes to some extent, none of them has been described as an essential gene for cell survival or a driver of tumorigenesis [30], [31], [32], [33], [34], [35]. From this list, we selected the cell surface receptor MPZL1 (*myelin protein zero-like 1*), as at least some research on this protein has so far been linked to its functional role in cancer [33], [36], [37].

Next, we investigated the expression change of MPZL1 in relation to its genomic status. First, when correlating the *MPZL1* CNV status and with its mRNA expression levels, we observed a progressive mRNA elevation from diploid to amplified cancers across different tumor types (Figure 1G, Suppl. Figures S1C and F). Next, we investigated MPZL1 in a unique patient cohort for which we had access to the genomic status, mRNA expression, and protein levels (Figure 1H). Specimens were classified into “normal liver”, “diploid HCC” or “MPZL1 gain HCC” via comparative genomic hybridization (CGH). Subsequent mRNA microarray and Western blot analyses revealed a marked increase of MPZL1 mRNA expression (Figure 1I), as well as MPZL1 protein expression in gained HCCs compared to normal liver and diploid HCCs (Figure 1J). Thus, these data show that genomic amplifications in MPZL1 not only increase its transcriptional levels but also protein levels, further suggesting MPZL1 as a suitable target for therapeutic strategies.

### MPZL1 is expressed at the plasma cell membrane in a variety of human solid cancers

To gain more insights into MPZL1 protein expression, we first screened human liver cancer cell lines for their protein abundance of MPZL1 with a commercially available antibody. Western blot analysis of whole cell lysates (WCLs) revealed MPZL1 expression, ranging from approximately 34 kDa to 54 kDa (Suppl. Figure S2A, left panel), possibly due to its known glycosylation [38]. Accordingly, when protein lysates were subjected to deglycosylation via treatment with PNGase F, a single band of around 29 kDa matching the estimated molecular weight of the native amino acids sequence of MPZL1 was revealed (Suppl. Figure S2A, right panel), reinforcing the validity of this antibody. Densitometry-based analysis of protein bands provided quantitative measurements of MPZL1 levels consistent with the band intensities observed for untreated samples. Moreover, publicly available CNV status data for *MPZL1* (cancer cell line encyclopedia – CCLE) showed a good correlation with protein expression levels (Suppl. Figure S2A), further demonstrating a causal relationship of amplification with high expression. Similarly, MPZL1 protein expression in lung and breast cancer cell lines also exhibited different patterns of glycosylation (Suppl. Figure S2B). However, while protein expression and CNV status correlated well for breast cancer, this was not always evident for lung cancer cell lines, possibly due to a lower protein turnover or mutations in related pathways. Additionally, a leukemia cell line (Molm13) with very low-to-absent MPZL1 expression was included as a negative control (Suppl. Figure S2B).

Based on protein levels we categorized cell lines as MPZL1-high or -low and established isogenic systems to study the biological relevance of MPZL1. MPZL1^high^ cell lines (Hep3B, SNU387, JIMT-1, and NCI-H2087) were subjected to CRISPR/Cas9-mediated knockout of *MPZL1*, whereas human *MPZL1* cDNA was overexpressed in MPZL1^low^ cell lines (Huh7 and HLE) and western blot analysis of whole cell lysates corroborated efficiency of genetic modifications (Suppl. Figure S2C). Interestingly, neither colony formation assays nor short-term cell proliferation assays showed differences in growth capabilities upon MPZL1 perturbation (Suppl. Figures S2D and E), supporting its potential role as bystander alteration in tumorigenesis and as a non-essential gene for cell survival.

In order to analyze the subcellular localization of MPZL1, we performed cell fractionation assays (Figure 2A), followed by Western blot analysis. These data revealed presence of MPZL1 protein in all membranous structures, with a preferential localization at the plasma membrane in both MPZL1^low^ (Figure 2B) and MPZL1^high^ cells (Figure 2C), further reassuring its value as a promising candidate for a CSP-targeted therapy. Next, we evaluated the expression of MPZL1 protein by immunohistochemistry in paraffin embedded human tissues of different cancer entities and normal tissues (Table 3). 2244 resected cancer samples across six different cancer entities as well as thirty normal healthy tissues (3 independent samples for each tissue) were included in the study. Histological examination revealed that majority of cancer tissues exhibited specific MPZL1 staining at the plasma membrane of cancer cells, whereas respective healthy tissues showed no specific staining or only faint, primarily intracellular MPZL1 expression (Figure 2D). Notably, normal tissues from other, not paired but essential tissue types (muscle, heart, bone marrow, brain, etc.), were clearly negative for MPZL1 (Figure 2D). Subsequently, we developed a scoring system for MPZL1 expression levels at the plasma membrane, analogous to the well-established scoring system for HER2 in breast cancer [39] (Figure 2E). For the majority of included cancer types, a high percentage of cases presented a high score (score 2 or 3), such as 47.9% of HCC (hepatocellular carcinoma), 39.6% of PDAC (pancreatic ductal adenocarcinoma) or 64.7% of CRC (colorectal cancer) samples (Figure 2F). Strikingly, 89% of TNBC (triple-negative breast cancer) cases presented a high score, with 66% and 23% of samples receiving scores 3 and 2, respectively. However, only 15.5% of conventional BRCA (breast cancer) samples showed high scores (score 2 or 3). In contrast, most normal tissues presented a score of 0, indicating no detectable MPZL1 expression, whereas some tissues (e.g. liver, stomach, and colon) revealed low expression (score 1). Only the uterus and testis presented a considerable higher score (score 2) (Figure 2G). Of note, myelinated tissues (brain, nerves etc.) were completely negative for MPZL1, although its name would be suggestive for positivity in these tissues. Therefore, these results not only reveal that MPZL1 is expressed at the plasma membrane in human cancer cells and cancer tissues but also demonstrate that high MPZL1 expression is restricted to certain cancer subgroups, whereas normal tissues have low expression, highlighting its therapeutic potential.

**Figure 2.**
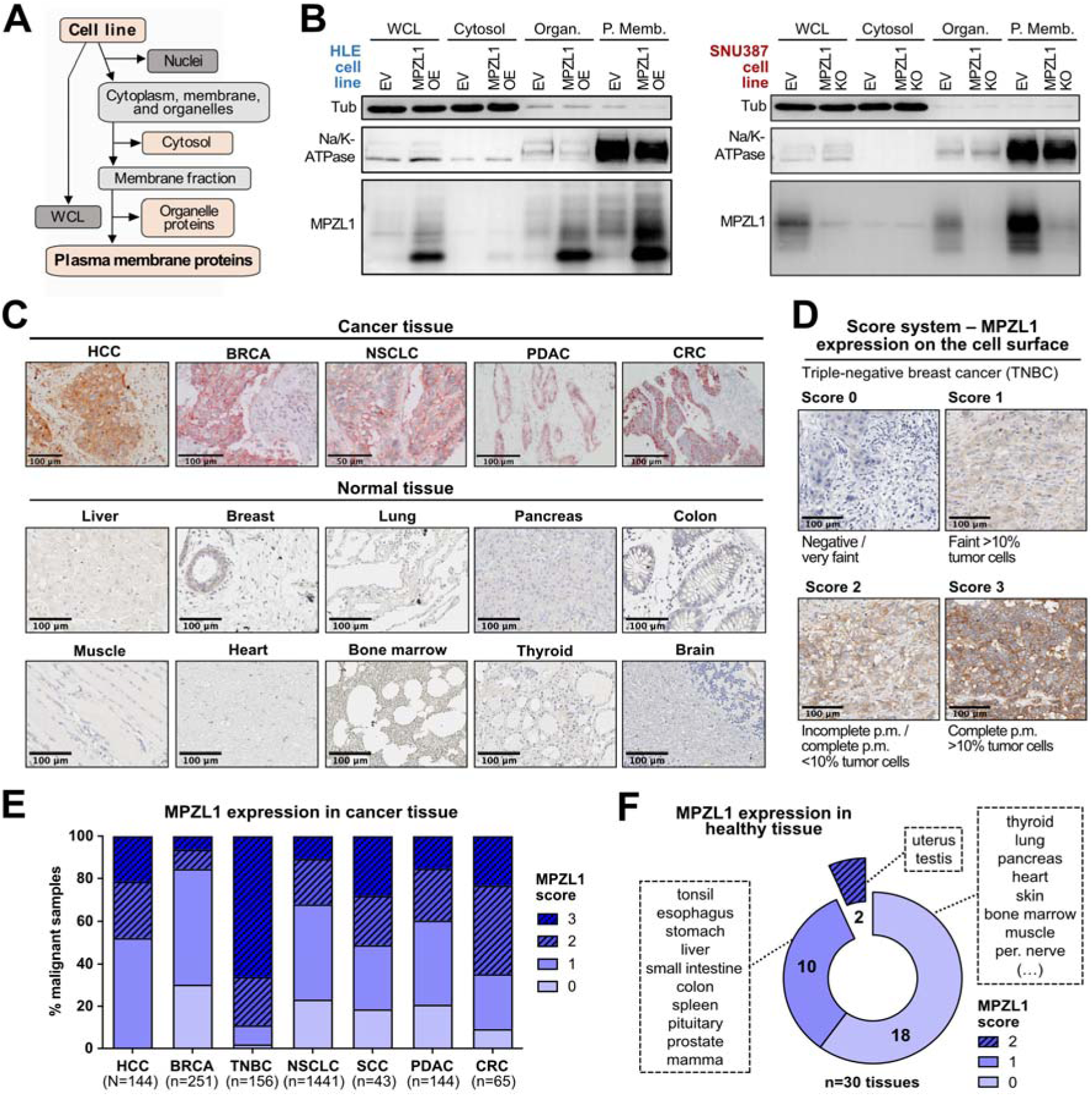
MPZL1 is expressed at the plasma membrane in a variety of human solid cancers. **(A) – (B) Cell fractionation assay. (A)** Schematic depicting the assay workflow and the resulting cellular fractions. **(B)** Immunoblotting analysis of MPZL1 levels on the different cellular fractions in isogenic liver cancer cell lines: MPZL1^low^ (HLE – left panel) and MPZL1^high^ (SNU387 – right panel). From left to right: whole cell lysate, cytosol, organelle proteins and plasma membrane proteins. Tubulin and Na/K-ATPase are used as markers of the cytosolic and plasma membrane fractions, respectively. *MPZL1* knock-out cells are generated with the MPZL1 targeting guide “sg1”. **(C) – (F) Immunohistochemistry.** Evaluation of MPZL1 expression at the protein level via IHC staining of human tissue microarrays (TMAs) using a commercially available antibody that binds to the intracellular region of MPZL1. For the cancer tissues, the following TMAs are utilized: HCC (n=144), BRCA (n=251), TNBC (n=156), NSCLC (n=1441), SCC (n=43), PDAC (n=144) and CRC (n=65), where HCC: Hepatocellular Carcinoma, BRCA: Breast Cancer, TNBC: Triple-Negative Breast Cancer, NSCLC: Non-Small Cell Lung Cancer, SCC: Squamous Cell Carcinoma, PDAC: Pancreatic Ductal Adenocarcinoma, and CRC: Colorectal Cancer. For the normal tissues, one microarray containing 30 different healthy tissues (3 independent patients each) is used: cerebrum cortex/white matter, cerebellum cortex, pituitary adenohypophysis /neurohypophysis, uterus, cervix, ovary, testis, prostate, mamma, skeletal muscle, inner lining cells, peripheral nerve, skin, bone marrow, salivary gland, tonsil, thyroid, parathyroid, thymus, lung, heart, esophagus, stomach, pancreas, liver, small intestine, colon, spleen, kidney, and adrenal cortex/medulla. All TMAs are gathered by the Institute of Pathology in Heidelberg – NCT Tissue Bank. Additional details regarding the IHC staining procedure are provided in the Methods section. **(C)** Representative images of MPZL1 IHC staining of cancer tissues (*top*) and healthy tissues (*bottom*) retrieved from the TMAs. Images are either acquired manually utilizing a microscope, or obtained from whole-slide scans at 20X. Visualization is performed using QuPath and ImageJ (scale bar: 50-100_μ_m). **(D)** Score system to assess the expression of MPZL1 protein at the cell surface. Representative images of TNBC tissues are assigned to each score (0 to 3). Slides are scanned at 20X (scale bar: 100_μ_m). **(E)** – **(F)** Classification of all tissue spots in the TMAs according to the score system in (D). **(E)** Cancer tissues, and **(F)** normal tissues – the numbers in the donut plot indicate the distribution of tissue types assigned to each score.

### Generation of a specific antibody targeting extracellular domains of MPZL1-protein

Having demonstrated that MPZL1 could be a promising target to deliver anti-cancer therapy, we further wanted to prove its suitability. To do so, this would require detection of MPZL1 protein at the extracellular space. However, for the above described experiments we utilized a commercially available MPZL1-targeted antibody, which binds to the cytoplasmatic domain of the human protein, specifically around Leu209, and thus make it unsuitable for further therapeutic studies. Thus, we pursued the development of antibodies that specifically target the extracellular domain of MPZL1. Several mice were immunized with the extracellular fragment of human MPZL1 protein (aa 36-165) and monoclonal antibodies (mAbs) were derived by hybridoma technology. Antibody supernatants were then tested via flow cytometry and suitable clones were subjected to two consecutive rounds of subcloning, identifying the most specific subclone for MPZL1-mAb. RNA from the selected hybridoma was sequenced, and subsequent bioinformatic analysis identified the sequences of the variable domain responsible for the specificity, both heavy (V_H_) and *kappa* light (V_L_) chains. These sequences were then cloned in frame in two independent human backbone vectors coding, respectively, for constant heavy (C_H_) and *kappa* light (C_L_) chains [40] to produce a recombinant chimeric antibody (MPZL1-ChAb) composed of a human constant domain and a mouse variable region (Figure 3A).

**Figure 3.**
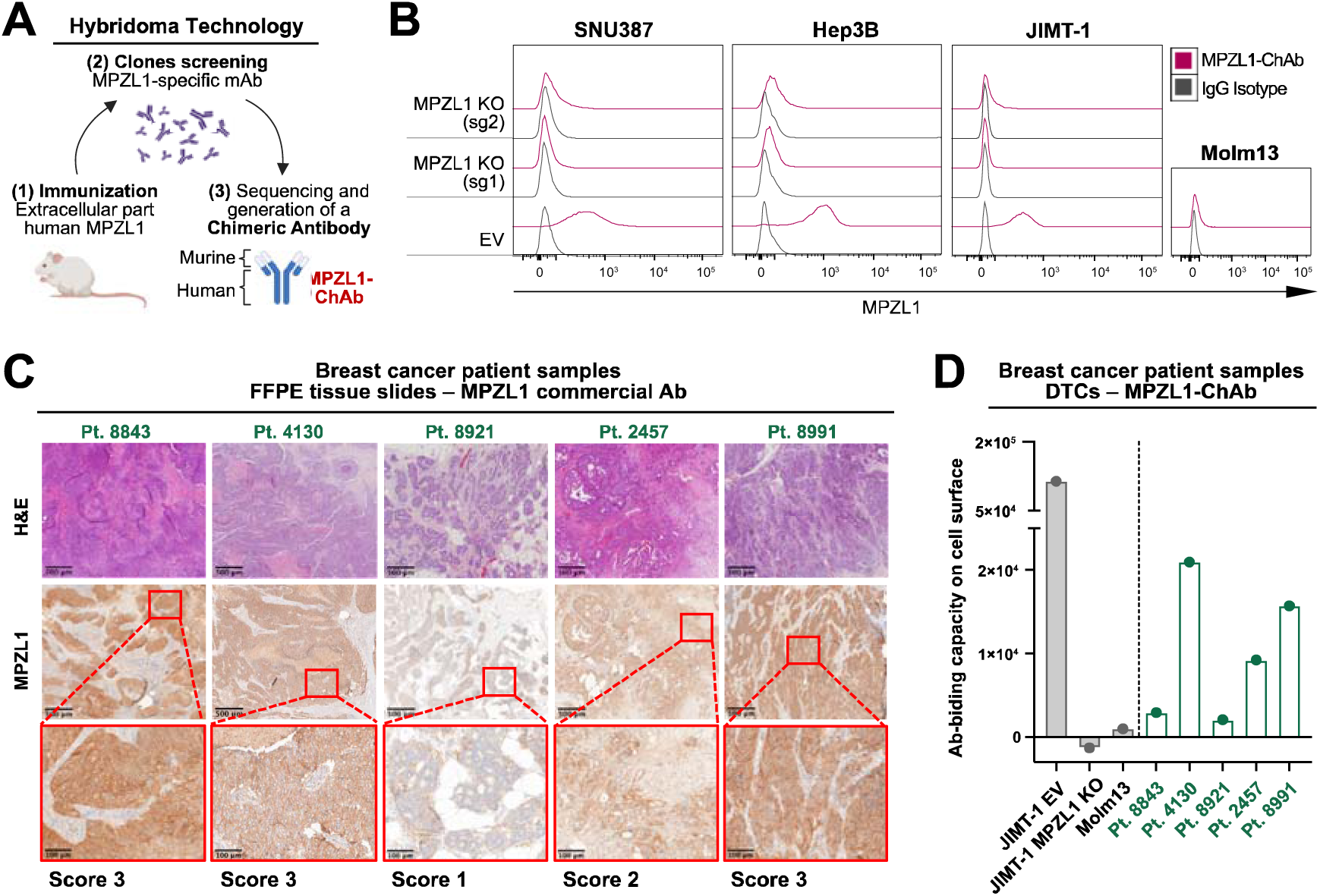
Generation of a specific antibody targeting the extracellular domain of MPZL1 protein. **(A)** Schematic of the experimental design. Several mice are immunized with the extracellular fragment of human MPZL1 protein, and monoclonal antibodies are derived using the hybridoma technology. The variable regions of the most specific antibody are then sequenced and cloned in a human backbone vector to generate a chimeric antibody (MPZL1-ChAb). Created with *BioRender.com*. **(B)** Histograms depict FACS data showing the specificity of MPZL1-ChAb by comparing its binding to MPZL1^high^ cell lines (EV) from different cancer types with their corresponding *MPZL1* KO cell lines (two independent guides used). Molm13 cells are used as a negative control, with absent endogenous MPZL1 expression. Cells are detached with *Versene*, and staining is performed by a first incubation with 100 µg/mL of either MPZL1-ChAb or IgG isotype, followed by a second incubation with a fluorochrome-coupled secondary anti-human antibody (1:200 dilution). **(C) – (C) Specificity of MPZL1-ChAb in breast cancer patient samples.** Matched dissociated tumor cells (DTCs) and FFPE tissue blocks from 5 independent breast cancer patients are acquired from an external vendor. **(C)** Representative images of H&E and MPZL1 IHC staining of FFPE-derived tissue slides. MPZL1 IHC staining is done with the commercially available antibody. Specific MPZL1 scores are assigned to the different patients according to the score system described in Figure 2D. Slides are scanned at 20X, and the images are visualized using QuPath and ImageJ, with two magnifications shown (scale bars: 500_μ_m and 100_μ_m). **(D)** Quantification of MPZL1-ChAb biding capacity (ABC) on the cell surface on the DTCs. Both MPZL1-ChAb and IgG isotype are directly labeled to a fluorochrome, and the binding specificity is analyzed via FACS. JIMT-1 EV, JIMT-1 MPZL1 KO and Molm13 cell lines are used as experimental controls. Additional details regarding the procedure are explained in the Methods section. In (B) and (D), data acquisition is done using the BD LSRFortessa flow cytometer (BD, Germany) and the BD FACS Diva software v8.0.1m, and analysis is carried out with FlowJo v10.

Given the non-essential nature of MPZL1, the MPZL1-ChAb was not expected to act as a therapeutic antibody but rather to serve as a carrier for therapeutic agents (antibody-drug conjugates, CAR-T cells, etc.), facilitating its specific delivery to target cells. To confirm this, the previously generated isogenic liver cancer cell lines were treated with increasing concentrations of MPZL1-ChAb. Indeed, regardless of their MPZL1 status, no differences in cell survival were observed between the different cell line pairs (Suppl. Figure S3A).

Subsequent validation of MPZL1-ChAb binding specificity was performed through flow cytometry experiments using isogenic cell lines without permeabilization to ensure that the observed binding occurs solely through the extracellular fraction on the cell surface. Using MPZL1^high^ cell lines from cancer entities of interest (HCC, BRCA, LUAD) and their isogenic counterparts with *MPZL1* KO (two independent sgRNAs), we observed specific binding of MPZL1-ChAb in all MPZL1^high^ cell lines compared to the IgG isotype control, whereas no binding was detected in the cell lines with *MPZL1* KO. Moreover, the antibody did not bind to Molm13 leukemia cells, which are negative for MPZL1 (Figure 3B). We also investigated its potential cross-reactivity with the murine MPZL1 protein using primary mouse liver cancer cell lines. While no binding of MPZL1-ChAb was detected in these cell lines, antibody binding was re-established when human *MPZL1* cDNA was introduced in murine cells (Suppl. Figure S3B), corroborating the specificity of MPZL1-ChAb for human MPZL1 protein. However, we found that MPZL1-ChAb was not suitable for other assays, such as immunohistochemistry or Western blot, potentially due to the recognition of conformational or discontinuous epitopes, which can be lost upon protein denaturation during fixation [41].

As our new generated MPZL1-ChAb was not suitable for immunohistochemistry it would be important for prospective clinical settings to ensure that its binding capacity to tumors cells is closely reflected by the binding capacity of the commercially available MPZL1 antibody we used for immunohistochemistry of human tumors. To address this we obtained matched sets of dissociated tumor cells (DTCs) – cryopreserved viable single cell suspensions [42] – and FFPE tissue blocks from five different breast cancer patients, who underwent tumor resection. Upon immunohistochemical staining with the MPZL1 commercial antibody, these five tissues were classified according to their MPZL1 expression employing the scoring system introduced above. We identified tissues with either high or low expression, including score 3 (three patients), 2 and 1 (one other patient each) (Figure 3C). Additionally, the extracellular expression of MPZL1 on matching DTCs was analyzed using the newly developed MPZL1-ChAb, directly labeled to a fluorochrome. Quantification was performed by measuring the antibody binding capacity (ABC), which represents the amount of antibodies a cell can bind and correlates with the amount of specific antigens expressed on the cell surface. While tumor cells of patients 4130 and 8991 (MPZL1 IHC score 3) exhibited a high ABC, tumor cells of patient 2457 (MPZL1 IHC score 2) displayed a distinctly lower ABC value, and tumor cells of patient 8921 (MPZL1 IHC score 1) barely bound the antibody. An outlier was patient 8843 (MPZL1 IHC score 3), who did not show strong binding of MPZL1-ChAb (Figure 3D).

In summary, these results demonstrate the specificity of our new developed MPZL1-ChAb towards human MPZL1 protein and suggest that the MPZL1 commercial antibody can be used to screen patients suitable for MPZL1-targeted therapies.

### Generation of anti-MPZL1 CAR-T cells as a therapeutic approach for MPZL1 expressing cancers

In order to develop a MPZL1-specific therapy we chose a CAR-T cell approach, as they have the ability to directly target cells that express the specific antigen for which they are designed on their surface [43]. Using sequences of the variable heavy (V_H_) and *kappa* light (V_L_) chains of the MPZL1-ChAb we built specific scFvs, which were introduced in a second-generation CAR composed of both stimulatory (CD3ζ) and co-stimulatory (CD28) signals. Additionally, the construct is linked to a P2A sequence to induce co-expression of a reporter gene (LNGFR). These components were incorporated in a SFGγ retroviral vector, which was ultimately used to generate retroviral supernatants employed for the transduction of T cells (Figure 4A).

**Figure 4.**
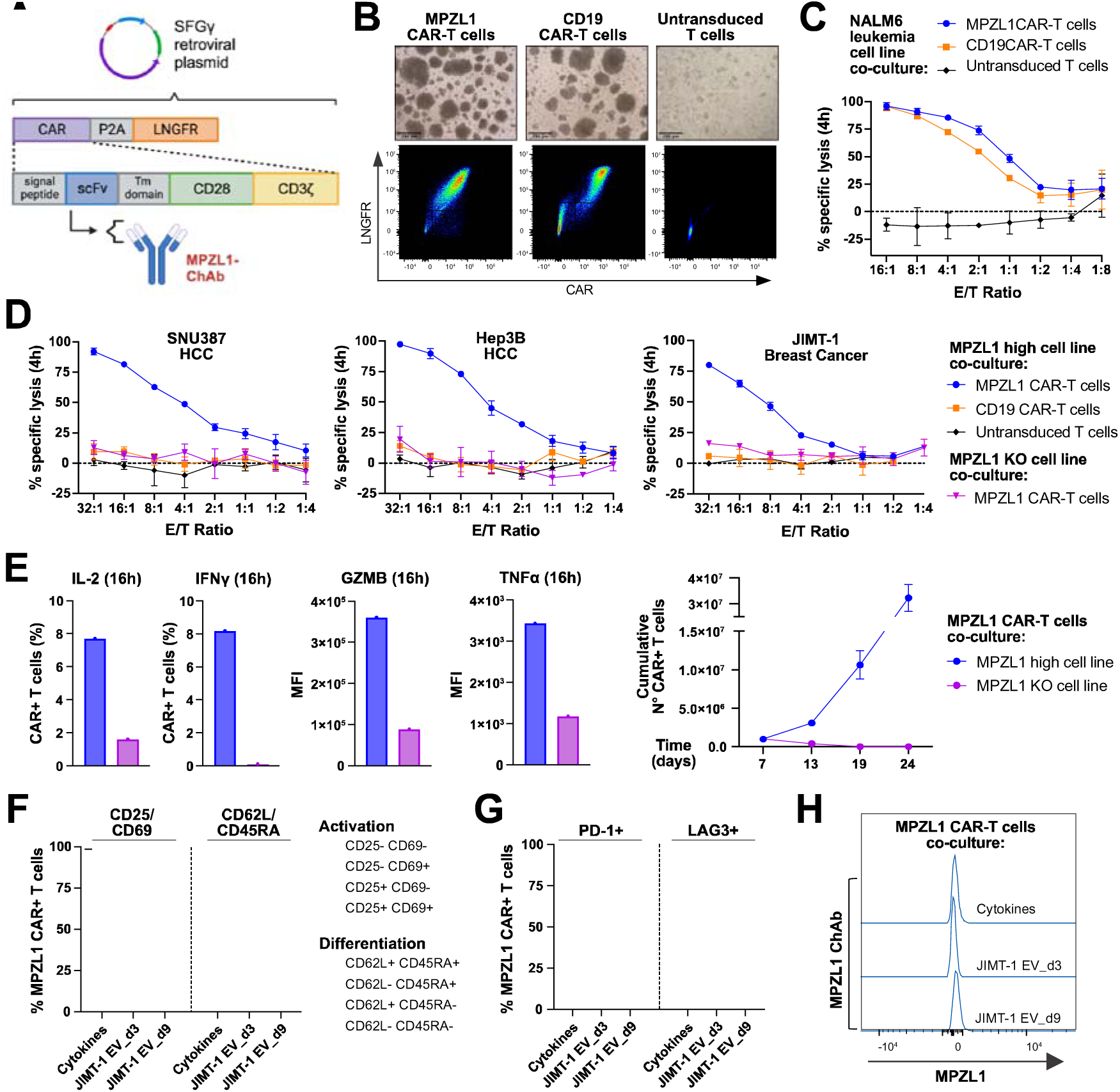
MPZL1 CAR-T cells exhibit high killing efficiency and specificity *in vitro*. **(A)** Schematic shows the structure of MPZL1 chimeric antigen receptor transgene. Created with *BioRender.com*. **(B)** T cells utilized 5 days after transduction. **Top panel**: representative bright-field (BF) microscopic images of MPZL1 CAR, CD19 CAR and untransduced T cells (scale bar: 250_μ_m). **Bottom panel**: dotplots show FACS data depicting the percentage of CAR+ T cells for each of the cell types. Staining is performed by a first incubation using AF647 anti-GAM F(ab’)_2_, followed by a blocking step with mouse serum, and a second incubation with PE-LNGFR antibody. **(C)** – **(D)** Bioluminescence-based cytotoxicity assay co-culture of the different T cells with cancer target cells during 4 hours at decreasing effector-to-target cell ratios. Cancer cells are stably transduced with a FFLuc-GFP construct. RPMI medium and 0.2% triton X-100 are used as negative and positive lysis controls, respectively. For the readout, luciferin substrate is added to the medium, and bioluminescence is detected in a microplate reader. **(C)** MPZL1 and CD19 CAR-T cells exhibit efficient lysis of NALM6 leukemia cells (MPZL1^high^; CD19^high^). **(D)** MPZL1 CAR-T cells demonstrate specific lysis of various MPZL1^high^ cancer cells lines (SNU387, Hep3B, JIMT-1) compared to their respective *MPZL1* KO cell lines. **(E)** – **(H)** Co-culture assays of MPZL1 CAR-T cells with JIMT-1 MPZL1^high^ or JIMT-1 *MPZL1* KO target cells. **(E) Left panel**: analysis of cytotoxic cytokine secretion by CAR-T cells. Co-cultures are conducted for 16 hours, with a cocktail of Brefeldin A and Monensin added during the last 4 hours to retain the cytokines within the cells. Bar plots show FACS data depicting the secretion of IL-2, IFN_γ_, GZMB, and TNF_α_ by MPZL1 CAR-T cells in the presence and absence of the specific antigen. **Right panel**: antigen-dependent proliferation assay. MPZL1 CAR-T cells are subjected to several consecutive antigen-specific stimulation rounds with respective target cells, which had been irradiated to supress proliferation. MPZL1 CAR-T cells are counted and replated with fresh target cells every six days. **(F)** – **(H)** FACS phenotypic analysis of CAR-T cells 22 days after transduction. Three independent groups of MPZL1 CAR-T cells are included: re-stimulated with JIMT-1 EV cells 3 and 9 days before analysis (respectively named “*JIMT-1 EV_d3*” and “*JIMT-1 EV_9*”) and non-specifically stimulated cells (“*Cytokines*”). **(F) Left part**: CAR-T cells arranged according to their activation pattern determined by the markers CD25 and CD69. **Right part**: CAR-T cells classified based on their differentiation pattern defined by CD62L and CD45RA markers. **(G)** Percentage of CAR T cells expressing the exhaustion markers PD-1 and LAG3. **(H)** Histograms show FACS data depicting binding of MPZL1-ChAb to the differently stimulated cell types, excluding potential trogocytosis. Staining is done by incubation with 100 µg/mL of MPZL1-ChAb, followed by incubation with a secondary anti-human antibody (1:200) coupled a fluorochrome. Further details about antibodies used and staining procedures followed for both intracellular cytokine analysis and phenotypic analysis can be found in the Methods section. In all cases, acquisition is performed using the Cytek^®^ Aurora (Cytek Biosciences, USA) and the SpectroFlow software, while analysis is carried out with FlowJo v10.

T cells were isolated from buffy coats of anonymous healthy donors, activated *in vitro* with CD3/CD28 beads and transduced with respective retroviral vectors. Untransduced T cells (UT T cells) and CD19 CAR-T cells served as experimental controls. Successful CAR expression on the cell surface was evaluated by extracellular staining with CAR and LNGFR antibodies via FACS analysis. Typically, five days after transduction, around 50-60% of CD19 and 80%-90% of MPZL1 CAR-T cells were CAR positive, while there was no CAR expression detected on UT T cells (Figure 4B).

Four distinct MPZL1 CAR constructs were designed and tested for their performance: two with V_L_+V_H_ (MPZL1.23 and MPZL1.44) and two with V_H_+V_L_ (MPZL1.24 and MPZL1.66) configurations (Suppl. Figure S4A). Staining of the different CAR-T cells with MPZL1-ChAb did not reveal any binding in the FACS analysis, thereby excluding the possibility of T cell fratricide [44] (Suppl. Figure S4B). Moreover, T cells transduced with the different MPZL1 CAR constructs exhibited comparable killing capacity of MPZL1^high^ breast cancer cells (Suppl. Figure S4C). When we evaluated proliferation, viability, and CAR expression progression over time we found while CAR expression was similar for the four designs, the configuration V_L_+V_H_ achieved 85-90% of CAR+ cells already on day 4 after transduction and stayed constant until day 13, whereas CAR expression started to decline by day 9 for V_H_+V_L_ constructs. Additionally, MPZL1.23 and MPZL1.44 CAR-T cells restarted proliferating 6 days upon transduction (3-fold from day 6 to 9), with a sustained viability of 60-70%, whereas MPZL1.24 and MPZL1.66 CAR-T cells did not grow and started dying by day 13, as confirmed by their viability of only around 20% at day 17. Finally, although CD19 CAR-T cells grew roughly 2-fold faster than MPZL1.23/44 CAR-T cells (data not shown), the viability was similar in both cases (Suppl. Figure S4D). Phenotypical profiling did not reveal meaningful differences among the different CAR-T cells in CD4/CD8 (Suppl. Figure S4E) or CD62L/CD45RA subpopulations, which were in all cases mostly in a naïve status represented by CD62L+CD45RA+, as expected for unstimulated cells [45], [46], [47] (Suppl. Figure S4F). However, PD-1 expression levels were notably higher for MPZL1.24 and MPZL1.66 (70% and 66%, respectively) in comparison to MPZL1.23 and MPZL1.44 (30% and 50%, respectively) CAR-T cells (Suppl. Figure S4G). Consequently, we concluded that MPZL1 CAR-T cells with V_L_+V_H_ outperformed those with V_H_+V_L_ chains, and chose MPZL1.44 CAR-T cells (from now on referred to as just MPZL1 CAR-T cells) for further functional assays.

### MPZL1 CAR-T cells show high efficiency and specificity *in vitro*

To test killing capability of MPZL1 CAR-T cells, CAR-T cells were co-cultured with luciferase-expressing target cells over time and killing was assessed via bioluminescence-based cytotoxicity assays. NALM6 leukemia cells, which express both MPZL1 and CD19 antigens (Suppl. Figure S5A), exhibited up to 100% lysis in the presence of MPZL1 CAR-T cells or CD19 CAR-T cells (Figure 4C). Next, we utilized MPZL1^high^ cell lines from different cancer types including HCC, breast, lung, pancreas and glioblastoma (Suppl. Figure S5B) in this assay and found that MPZL1 CAR-T cells exhibited a robust killing capacity against all cancer cells, whereas CD19 CAR-T cells or UT T cells did not (Suppl. Figure S5C). Thus, regardless of the origin of the cancer entity MPZL1 CAR-T cells can execute their antitumor function.

In order to test the antigen-specific killing capacity of MPZL1 CAR-T cells we used isogenic pairs of MPZL1^high^ cancer cell lines with *MPZL1* KO (Suppl. Figure S2C) in co-culture assays. In 4-hour killing assays, MPZL1 CAR-T cells exhibited 80% to 100% lysis of all MPZL1^high^ target cell lines at the highest E/T ratios utilized (blue curve). In contrast, we did not observe specific killing of respective isogenic *MPZL1* KO cell line pairs (purple curve), as the lysis curves were comparable to those induced by CD19 CAR-T cells or UT T cells (Figure 4D). Furthermore, we performed a short-term stimulation assay to evaluate the release of cytotoxic cytokines by MPZL1 CAR-T cells upon exposure to the specific antigen. Whereas no IFNγ was detected in the absence of the MPZL1 target antigen, 8% of MPZL1 CAR+ T cells released this cytokine upon specific stimulation. Similarly, we observed a 4-fold increase in the percentage of cells producing IL-2 (from 1.63% to 7.71%), as well as a 4-fold increase in MFI for GZMB and a 3-fold increase for TNFα when MPZL1 CAR-T cells were stimulated with MPZL1^high^ target cells (JIMT-1) compared to respective *MPZL1* KO target cells (Figure 4E, left panel). Finally, we conducted a long-term stimulation assay to evaluate the proliferative response of MPZL1 CAR+ T cells in the presence of the corresponding antigen over an extended period of time. In the co-culture of MPZL1 CAR-T cells with MPZL1^high^ target cells (JIMT-1), we detected a 3-fold increase in the cumulative amount of MPZL1 CAR+ T cells across successive re-stimulation rounds (from 1M to 3M to 10.5M to 32.3M). Conversely, when co-cultured with respective *MPZL1* KO cells, the same CAR+ T cells underwent a 2-fold decrease in the total cell count after the first six-day interval, resulting in rapid cell death (Figure 4E, right panel).

Next, phenotypical profiling of MPZL1 CAR-T cells at a late time point (day 22 after transduction) was performed, including three independent groups: CAR-T cells re-stimulated with MPZL1^high^ JIMT-1 cells 3 and 9 days before analysis (respectively named “*JIMT-1 EV_d3*” and “*JIMT-1 EV_d9*”) and non-stimulated CAR-T cells, cultured with cytokines (“*Cytokines*”). While antigen-stimulated CAR-T cells were principally CD25+CD69-/+, indicating a late activation state, 12% of unstimulated CAR-T cells remained CD25-CD69-, suggesting a still naïve status. Similarly, unstimulated MPZL1 CAR-T cells were mostly CD62L+CD45RA+ (42.3%), pointing again to a primarily naïve status, with a limited presence of differentiated cells (CD62L+/-CD45RA-) (30% both phenotypes combined). Strikingly, upon antigen-specific stimulation, MPZL1 CAR-T cells transitioned to CD62L+CD45RA-(47.2% on “d3” and 18.7% on “d9”) and CD62L-CD45RA-(46.1% on “d3” and 68.4% on “d9”) differentiation states, which indicates central memory and effector memory phenotypes, respectively [45], [46], [47]. Together, these two differentiated phenotypes constituted around 90% of total CAR+ T cells (Figure 4F). This finding was consistent with the previous observation at day 11 after transduction, where unstimulated CAR-T cells were mainly naïve (Suppl. Figure S4F). Interestingly, a high percentage of stimulated MPZL1 CAR-T cells presented the exhaustion markers PD-1 (76.9% on “d3” and 70.2% on “d9”) and LAG3 (47.8% on “d3” and 14.5% on “d9”), in contrast to CAR-T cells cultured with cytokines (25.5% PD-1+ and 9.9% LAG3+) (Figure 4G). The phenomenon of CAR-T cell exhaustion due to continued antigen stimulation can dampen their persistence and functionality, but in certain cases it becomes essential to avoid uncontrolled proliferation which can result in unwanted adverse effects [48]. Finally, when the same groups of MPZL1 CAR-T cells were stained with MPZL1-ChAb, no antibody binding was detected via FACS analysis (Figure 4H), confirming the absence of MPZL1 expression on the surface of CAR-T cells and thus excluding the possibility of trogocytosis, a phenomenon that can lead to CAR-T cell fratricide [49].

Taken together, these results demonstrate the ability and specificity of MPZL1 CAR-T cells to rapidly eliminate MPZL1+ target cells *in vitro* with a marked increase in cytotoxic cytokine production and accelerated proliferation when exposed to the specific antigen.

### MPZL1 CAR-T cells are highly efficient and specific to target MPZL1-expressing human xenograft tumors in vivo

As our *in vitro* experiments showed promising results, we next tried to test the effect of MPZL1 CAR-T cells *in vivo*. As a first approach we used a human HCC cell line-derived mouse xenograft model. HCC tumors were generated by subcutaneously injecting either Hep3B MPZL1^high^ cells or the respective isogenic Hep3B *MPZL1* KO cells in the flanks of immunodeficient NSG (NOD.Cg-Prkdc^scid^Il2rg^tm1Wjl^/SzJ) mice [50]. When tumors reached an average volume of 360 mm^3^, a single dose of 1*10^7^ MPZL1 CAR-T cells (right after primary *in vitro* expansion) was intravenously applied and tumor growth monitored over time (Figure 5A). Treatment with MPZL1 CAR-T cells provided a significant survival advantage of at least 20 days in mice harboring MPZL1^high^ tumors compared to those with *MPZL1* KO (Figure 5B). While mice having Hep3B *MPZL1* KO tumors did not respond to the treatment, those with Hep3B MPZL1^high^ tumors rapidly experienced tumor regression (Figure 5D). In relation to the time of CAR-T cell administration (day 0), Hep3B *MPZL1* KO tumors continued growing until the animals were sacrificed due to their large tumor volumes (Figure 5E, left panel). However, although Hep3B MPZL1^high^ tumors still increased in size by an average of 1.1-fold by day 4, this was followed by a 0.28-fold decrease already by day 8 (one fourth of the initial volume, p-value: <0.0001), therefore demonstrating a very quick and potent response to the therapy (Figure 5E, right panel). Strikingly, 7 out of the 11 MPZL1^high^ tumors were completely eradicated and did not grow back (follow-up until day 60), and only the remaining 4 tumors underwent regrowth (Figure 5D). Importantly, IHC staining confirmed the specific expression of MPZL1 on the cell surface of cancer cells within relapsed Hep3B MPZL1^high^ tumors, in contrast to Hep3B *MPZL1* KO tumors where MPZL1 expression was absent (Figure 5C). Additionally, we performed a similar experiment with human breast cancer tumors by injecting subcutaneously either JIMT-1 MPZL1^high^ cells or the respective JIMT-1 *MPZL1* KO cells in NSG mice. Again, MPZL1 CAR-T cell treatment provided a significant survival advantage of at least 17 days to mice harboring MPZL1^high^ tumors compared to those with *MPZL1* KO tumors (Suppl. Figure S6A). While JIMT-1 *MPZL1* KO tumors did not respond, JIMT-1 MPZL1^high^ tumors underwent noticeable tumor regression several days post-treatment (Suppl. Figure S6C). In this case, in relation to the time of CAR-T cell administration (day 0), JIMT-1 *MPZL1* KO tumors experienced a size increase of 2.16-fold on average already by day 9, and continued growing unstoppably until the animals had to be sacrificed (endpoint) (Suppl. Figure S6D, left panel). On the other hand, JIMT-1 MPZL1^high^ tumors exhibited on average a 1.33-fold increase in size by day 9, followed by a 0.61-fold decrease by day 19, and a subsequent recovery of nearly the initial tumor size by day 30 (Suppl. Figure S6D, right panel), resulting in evident tumor regrowth at the experimental endpoint (day 44). The regression in size (p-value: 0.0065) of JIMT-1 MPZL1^high^ tumors observed between days 9 and 19 proved that a single dose of MPZL1 CAR-T cells sufficed to induce a specific therapeutic response *in vivo*. Similar to HCC tumors, IHC staining revealed precise cell surface expression of MPZL1 in the cancer cells of JIMT-1 MPZL1^high^ tumors, compared to the negative staining observed in JIMT-1 *MPZL1* KO tumors (Figure 5B).

**Figure 5.**
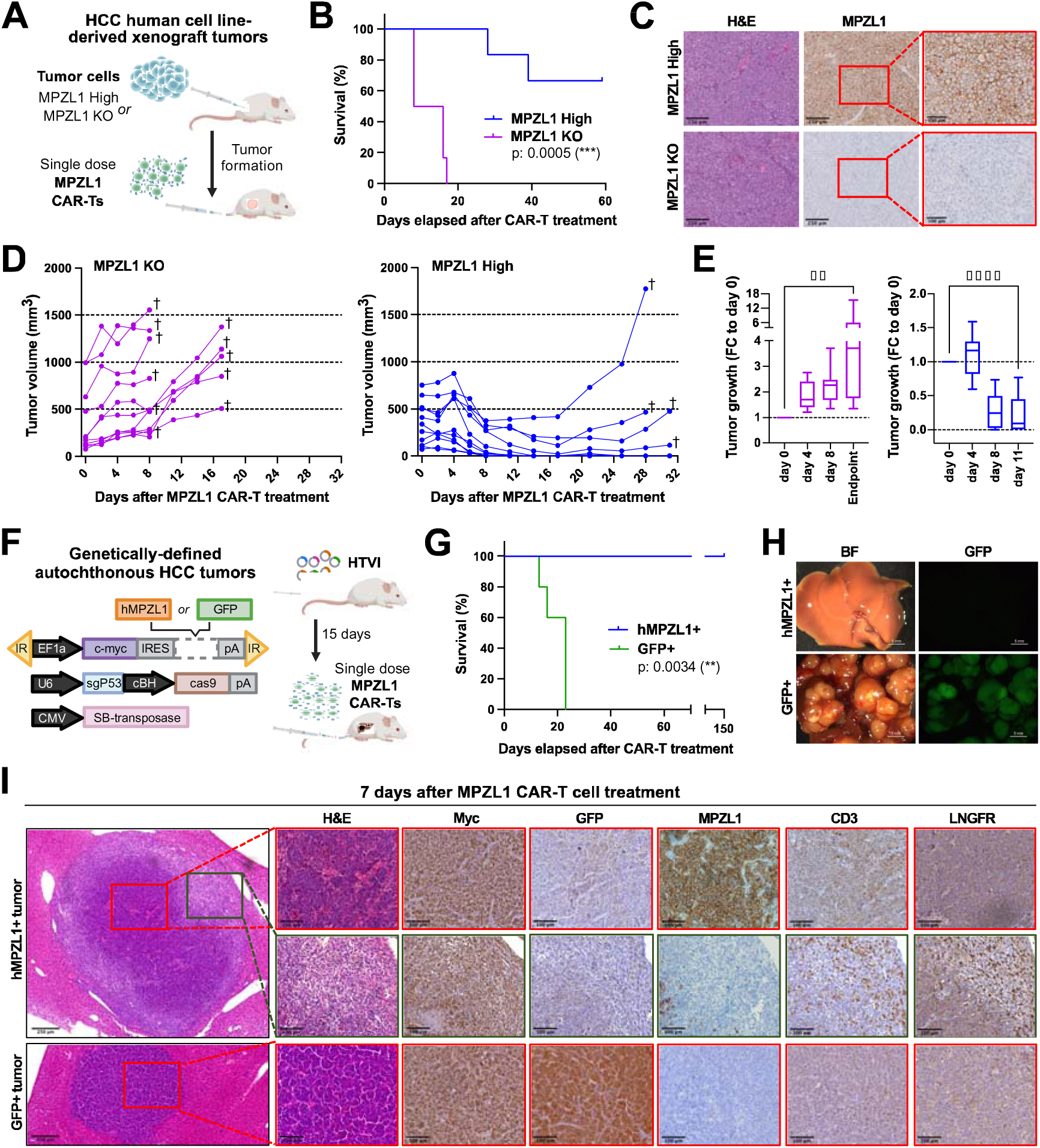
MPZL1 CAR-T cells efficiently and specifically target and eradicate MPZL1-expressing tumors in two distinct HCC mouse models. **(A) – (E) Cell line-derived xenograft tumor model**. **(A)** Schematic of the experimental setup: development of human HCC cell line-derived xenografts in NSG mice, followed by treatment with MPZL1 CAR-T cells. Created with *BioRender.com.* Tumor formation is induced by subcutaneous injection of either Hep3B EV (MPZL1^high^) or Hep3B *MPZL1* KO cells (5×10^6^ cells/flank). A single dose of 10×10^6^ MPZL1 CAR-T cells is applied intravenously when tumors reach an average volume of 360 mm^3^ (n=6 mice/group). **(B)** Kaplan-Meier curve shows survival analysis of the response to MPZL1 CAR-T treatment, with statistics derived from Log-rank test. **(C)** Representative images of FFPE tissue slides from both animal groups: H&E staining (scale bar: 250_μ_m) and MPZL1 IHC staining (two magnifications, scale bars: 250_μ_m and 100_μ_m). **(D)** Tumor growth curves display the volume progression (mm^3^) of *MPZL1* KO tumors (left panel) and MPZL1^high^ tumors (right panel) after therapy administration (day 0). **(E)** Quantification of the tumor growth curves in (D). Boxplots illustrate the variation in size of *MPZL1* KO tumors (left panel) and MPZL1^high^ tumors (right panel) at the selected time points relative to treatment administration (day 0) in terms of fold-change (FC), where values >1 represent an increase in tumor volume, whereas values <1 indicate a decrease thereof. *Endpoint* refers to the volume of each tumor at the time when the respective animal was sacrificed. Statistical analysis done via one-way ANOVA, and Tukey’s multiple comparisons test (**:p<0.01, ****:p<0.0001). **(F) – (I) Genetically-defined autochthonous HCC mouse model**. **(F)** Schematic of the experimental setup: induction of genotype-specific liver tumor development in NSG mice via hydrodynamic tail vein injection (HTVI), followed by treatment with MPZL1 CAR-T cells. Created with *BioRender.com*. Tumor formation is induced by rapid HTVI of a NaCl solution containing a plasmid cocktail, leading to overexpression of c-MYC, coupled with either human MPZL1 cDNA or GFP, and knockout of the p53 gene. The two groups are named as MycOE;p53-/-;MPZL1 (hMPZL1+) and MycOE;p53-/-;GFP (GFP+) (n=5 mice/group). A single dose of 10×10^6^ MPZL1 CAR-T cells is applied intravenously 15 days after HTVI. **(G)** Kaplan-Meier curve shows survival analysis of the response to MPZL1 CAR-T treatment, with statistics derived from Log-rank test. **(H)** Representative bright-field (BF) and fluorescent GFP stereomicroscopic images of resected livers (scale bar: 5mm). **(I)** Short-term experiment using the same plasmid configurations (hMPZL1+ and GFP+) for HTVI as shown in panel (F) (n=5 mice/group). A single dose of 10×10^6^ MPZL1 CAR-T cells is applied intravenously 30 days after HTVI, and mice are sacrificed 7 days following CAR-T therapy. H&E staining exhibits a complete tumor nodule from each group (scale bar: 250_μ_m). Selected regions are shown at higher magnification (scale bar: 100_μ_m) for both H&E and IHC stainings of target markers including Myc, GFP, MPZL1, CD3, and LNGFR. CAR-T cell infiltration is detected in hMPZL1+ tumors, while it is absent in GFP+ tumors. Additional details about antibodies and staining procedures can be found in the Methods section.

### MPZL1 CAR-T cells eradicate autochthonous HCC tumors in vivo

While human xenograft models provide valuable information to evaluate the CAR-T cell killing efficiency, subcutaneously implanted tumor cells do not resemble physiological tumor formation. Thus, we generated an autochthonous liver tumor model to test the impact of MPZL1 CAR-T cells on liver tumors in their native environment. Using hydrodynamic injection of plasmids encoding c-MYC coupled either to MPZL1 cDNA (MycOE;p53-/-;MPZL1) or GFP (MycOE;p53-/-;GFP) in conjunction with CRISPR/Cas9 mediated knockout of p53, we produced autochthonous liver tumors in NSG mice either with or without MPZL1 expression. A single intravenous dose of 1*10^7^ MPZL1 CAR-T cells (directly after primary *in vitro* expansion) was administered 15 days after HTVI and tumor growth was monitored over time (Figure 5F). While MycOE;p53-/-;GFP mice rapidly developed large tumors and had to be sacrificed by day 23 after treatment, MycOE;p53-/-;MPZL1 mice did not show any signs of tumor development and survived until the experimental endpoint (day 100) (Figure 5G). Indeed, MycOE;p53-/-;GFP mice showed multiple GFP-positive tumors, while no tumor nodules were found in MycOE;p53-/-;MPZL1mice at the end of the experiment (Figure 5H). To ensure that expression of MPZL1 per se had no detrimental effect to liver tumor formation we also generated a cohort of MycOE;p53-/-;GFP mice and MycOE;p53-/-;MPZL1 mice without CAR-T cell treatment and found that tumor onset and survival of both groups did not show any significant difference (Suppl. Figure S6E). Importantly, IHC staining of resulting MycOE;p53-/-;MPZL1 tumors revealed strong membranous expression of human MPZL1 protein on the cell surface of the tumor cells, whereas no signal was detected in MycOE;p53-/-;GFP tumors (Suppl. Figure S6F).

As our previous experiments solely examined end points after MPZL1 CAR-T injection, we wanted to understand the fate of tumors in early stages after CAR-T cell induction. Thus, we induced tumors via HTVI as described above (Figure 5F), injected mice received a single dose of 1*10^7^ MPZL1 CAR-T cells 4 weeks after HTVI and livers were harvested 7 days after the treatment. Microscopical examination of MycOE;p53-/-;GFP tumors revealed multiple tumor nodules with no significant immune cell infiltration. However, most of the detected MycOE;p53-/-;MPZL1 tumor nodules displayed a ring of immune cells surrounding the tumor and invading its inner core, while other hMPZL1+ tumor nodules were completely infiltrated by immune cells, with no histologically detectable tumor cells. Further immunohistochemical analyses of tumor sections revealed that MycOE;p53-/-;GFP tumors strongly expressed GFP and were negative for MPZL1, whereas MycOE;p53-/-;MPZL1 tumors were negative for GFP but showed a strong membranous expression of MPZL1 (Figure 5I). Characterization of infiltrating immune cells showed that the majority of immune cells in MycOE;p53-/-;MPZL1 were positive for CD3, and likely CAR-T cells as NSG mice do not have endogenous T cells. To further corroborate that these CD3+ cells were indeed CAR-T cells, we stained for LNGFR, a marker which is expressed from the CAR-construct. Consistently, CD3+ cells coexpressed LNGFR, confirming that they were MPZL1 CAR-T cells actively infiltrating the tumor core and eradicating MycOE;p53-/-;MPZL1 tumor nodules (Figure 5I).

In summary, these results demonstrate that MPZL1 CAR-T cells have high efficiency and specificity to completely eradicate MPZL1-positive tumors not only in human xenografts but also in autochthonous liver tumors *in vivo*.

## Discussion

The vast majority of therapeutic approaches developed in recent decades have been designed to target driver alterations responsible for tumorigenesis. However, identifying these drivers remains challenging [51] and most of these events can only be found in specific tumor types, hence only a small subset of patients can benefit from these new therapies [1], [11]. Here, we outlined an approach to exploit passenger amplifications, which are ubiquitous in solid cancers, as specific entry points for therapies, identified the cell surface protein MPZL1 as a promising candidate, and validated its utility for precision medicine.

Using a multiomics approach integrating publicly available chromosomal amplification status of human tumors, RNA expression of healthy and tumor tissue as well as annotated cell surface proteins, we were able to map potential CSPs, which are amenable to serving as entry points in cancer cells. Interestingly, we found that most of them were located in chromosome 1q (chr. 1q) and in lower number also on chromosome 8q (chr. 8q). Amplification of the long arm of chromosome 1 (chr. 1q) is one of the most frequent chromosomal aberrations detected in various solid cancers such as HCC, BRCA and LUAD [7]. Although several genes within this chromosomal region have been proposed to play an important role in the pathogenesis of HCC [52], only *MCL1* (*myeloid cell leukemia-1*), which encodes an intracellular antiapoptotic protein, has been defined as a driver gene that promotes cell survival by preventing apoptosis [24]. Accordingly, various inhibitors for MCL1 have been developed but targeting MCL1 protein remains challenging due to its role in important cellular functions as well as the emergence of compensatory mechanisms that lead to therapy resistance [53]. Interestingly, none of the annotated CSPs located on chr. 1q are defined as driver genes or described as being involved in HCC tumorigenesis [21], [24], [52]. However, although it would be potentially even more promising if a cell surface protein is a driver gene, we speculated that this would not be necessary for targeted therapy. It is also interesting to speculate that designing a therapeutic entry point, which targets more than one amplified gene in one or more chromosomal region (e.g. chr. 1q and chr. 8q) could lead to higher specificity but this approach would certainly also diminish the potential patient group, which could be included in such an approach. Anyhow, this approach can be used in the future to broaden the spectrum of further therapeutic targets.

Our approach identified *myelin protein zero-like 1* (*MPZL1*), a glycosylated cell surface receptor belonging to the immunoglobulin superfamily [38], whose role in promoting migratory and invasive capabilities in various tumor types has been described in several publications [33], [36], [37]. Importantly, our immunohistochemistry results for MPZL1 expression showed a striking difference between tumor and corresponding healthy tissues, and confirmed MPZL1 cell surface expression in a high percentage of patients across cancer types. Notably, 89% of triple-negative breast cancer (TNBC) samples obtained a score of 2 or 3, compared to only 15.5% of general breast cancer samples. TNBC is currently the most aggressive subtype of breast cancer, lacking common biomarkers (ER, PR, HER2) and an effective targeted-therapy, which normally leads to an unfavorable prognosis [55]. Thus, this challenging cancer subtype could especially benefit from a therapy targeted to cancer cells by means of MPZL1. Even though MPZL1 expression is markedly lower in normal tissues compared to cancerous tissues –with only the reproductive tract showing a higher score– several other tissues, such as those forming the gastrointestinal tract (esophagus, stomach, intestine, colon), received a score of 1, indicating faint MPZL1 expression and potentially limiting its therapeutic window. However, the fact that low expression of a tumor associated antigen (TAA) is present in normal tissues does not necessarily indicate that it is not suitable for therapy. For instance, the claudin 18.2 (*CLDN18.2*) TAA has emerged as a promising target for the treatment of gastric and gastroesophageal junction cancers in recent years as it is often overexpressed in these tumors due to epithelial transformation and the loss of tight junction integrity [56], [57]. The claudin 18.2-targeting monoclonal antibody zolbetuximab has recently been approved by the FDA for the treatment of a specific subset of gastric cancer patients [58], and various other therapeutic strategies, such as claudin 18.2 antibody-drug conjugates and CAR-T cells, are already under advanced clinical trials [59], [60]. Notably, claudin 18.2 expression is detected in human normal gastric mucosa and respiratory system according to the human protein atlas [61]. Another example is provided by the HER2 TAA, whose dysregulation contributes to the development of various tumors, including breast, gastric, and endometrial cancers. In this case, the amplification of *ErbB2* gene, resulting in overexpression of HER2 protein at the cell surface, is the most common mechanism driving carcinogenesis [62]. Targeting HER2 with the mAb Trastuzumab has showed effectiveness against breast cancer without major adverse effects [9]. Similarly to claudin 18.2, HER2 expression is not restricted to tumors, as it is also present in several normal tissues, including male and female reproductive organs, lymphoid tissues, kidneys and the respiratory system, among others. [61]. Taken together, as TAAs are not only found in cancer tissues, it is key to balance their antitumor function with the risk of side effects in healthy tissues. These considerations provide a rational basis for exploring MPZL1-targeted therapies.

As there did not exist any antibodies targeting the extracellular region of MPZL1, we developed a highly specific and sensitive antibody against its extracellular domain. A major limitation is that this antibody (MPZL1-ChAb) is not suitable for applications other than flow-cytometry and thus cannot be used to stratify patients via immunohistochemistry scoring. This lack of functionality in other techniques may be due to its recognition of conformational or discontinuous epitopes [41]. Additionally, as MPZL1 is a highly glycosylated protein, we cannot exclude the possibility that MPZL1-ChAb recognizes a specific glycosylation pattern, which could affect binding when the protein is not in its native state, or even that it targets a tumor-specific glycopattern, as it has been reported for various antibodies in the literature [63]. Anyhow, using paired FFPE tumor tissue slides and dissociated tumor cells, we found that the commercially available MPZL1 antibody, which in our hands works excellent for immunohistochemistry, reliably mirrors the MPZL1 levels detected by our newly established antibody. Thus, we anticipate that the commercial MPZL1 antibody will serve as an ideal companion diagnostic tool to stratify patients who are likely to benefit from MPZL1-targeted therapies in future trials.

As a therapeutic approach, we chose CAR-T cells directed to MPZL1 and found highly specific cancer cell killing, along with significantly increased proliferation and cytotoxic cytokine secretion profiles upon antigen stimulation *in vitro*. Results from cell line-derived xenograft mouse models showed the capability of systemically injected MPZL1 CAR-T cells to reach tumors *in vivo*, leading to significant tumor size regression, or even elimination, of tumors expressing MPZL1 antigen. The fact that the anti-tumor response did not start directly upon treatment, but rather with some days delay, suggests that MPZL1 CAR-T cells can proliferate within the animal. Additionally, the persistence of MPZL1 expression in both Hep3B and JIMT-1 MPZL1^high^ tumors, which had undergone regrowth after an initial response to the treatment, excludes the possibility of an escape mechanism driven by antigen downregulation but rather implies that multiple doses could lead to a more sustained anti-tumor effect. Although subcutaneous xenograft models are the standard for preclinical studies regarding CAR-T cells [64], subcutaneously implanted tumor cells do not resemble physiological tumor formation. Thus, we complemented our in vivo studies with tumor induction by HTVI, which offers the advantage of generating tumors in the organ of origin and provides a more accurate representation of the tumor microenvironment. Strikingly, a single dose of MPZL1 CAR-T cells resulted in complete eradication of autochthonous liver tumors overexpressing human MPZL1 protein. Additionally, our short-term study confirmed the capability of MPZL1 CAR-T cells to infiltrate only those tumor nodules positive for MPZL1 antigen. However, one has to take into account that tumor formation was induced in immunocompromised mice (in both the xenograft and the HTVI models) and therefore the complexity of other immune cells is missing in this scenario. This needs to be considered for future clinical testing, in particular given the current knowledge that treating solid cancers with CAR-T cells presents a major challenge [65], [66]. Altogether, our results highlight the robust functionality of MPZL1 CAR-T cells in terms of specificity, tumor infiltration, and killing efficiency.

The assessment of on-target off-tumor toxicities [67] is the main limitation of our *in vivo* studies, as the scFv utilized to generate MPZL1 CAR-T cells recognizes human but not murine MPZL1 protein. We attempted to generate an antibody with dual human and murine MPZL1 specificities, but it was not successful. Building on the previous examples, despite expression of claudin 18.2 in some normal tissues, tolerable safety profiles have been reported in a Phase I clinical trial using second generation claudin 18.2 CAR-T cells for treatment of heavily pretreated, advanced gastric cancer patients [60], with several other early-phase clinical trials based on different CAR constructs currently ongoing [57]. Thus, dim target expression does not mean adverse reactions provided the expression in the tumor is significantly stronger. Considering our results, we speculate that the elevated expression of MPZL1 in tumor cells provides a therapeutic window, making the treatment less detrimental to healthy tissues with absent or only low expression of the target.

In summary, we demonstrate that bystander amplifications of cell surface protein-coding genes within large genomic amplifications can serve as Trojan horses for the specific targeting of cancer cells. This approach opens a new avenue for the development of therapeutic strategies to tackle solid cancers harboring specific amplifications, regardless of their origin. Thus, our findings constitute an innovative contribution to the field of anticancer drug development.

## Acknowledgements

We thank all members of the Tschaharganeh lab for constructive discussion and feedback during the project. We thank the German Cancer Research Center Central Animal Laboratory, the Center for Model Systems and Comparative Pathology of the Institute of Pathology Heidelberg, and the Tissue Bank of the National Center for Tumor Diseases for excellent technical support. This work was funded by ERC Starting Grant “CrispSCNAs” (grant number: 948172) awarded to DFT and ERC Starting Grant “CARsen” (grant number: 949667) awarded to JF) by the European Research Council. DFT is recipient of a grant from the Deutsche Forschungsgemeinschaft (DFG) (TS 293/3-1). Part of this work was performed within the DKFZ-Bayer alliance on cancer.

## Methods

### Integrative *in silico* approach for CSP target identification

Human cell surface protein-coding genes, encoding the respective cell surface proteins (CSPs) annotated in the CSPA (N=1492) [21], were surveyed for their genomic status in a publicly available pan-cancer dataset (ICGC/TCGA) comprising copy number variation (CNV) data from 2703 patients [22]. The analysis was done via the online platform cBioPortal, using the Onco Query Language “GAIN AMP” [68]. This tool employs GISTIC 2.0, an algorithm that follows a statistical approach to identify significant copy-number alterations in cancer genomes [69], assigning the following scores: -2 (homozygous deletion), -1 (heterozygous deletion), 0 (diploid state), 1 (copy-number gain) or 2 (high-level amplification). We considered amplified genes as those with assigned scores of 1 or 2. The chromosomal locations of the hits were retrieved using the *BioMart data mining tool* from Ensembl [70]. To identify tumor types with recurrent amplifications of chromosome 1q, publicly accessible copy number variation data from all cancer entities available at TCGA Firehose Legacy was explored with the Integrative Genomics Viewer Software (IGV) [23], [71]. For cancer types with frequent chr. 1q amplification, the CNV status of the identified hits was surveyed in the corresponding datasets (HCC, n=370; BRCA, n=1080; LUAD, n=516) via cBioPortal [24], [25], [26]. Next, differential expression analysis was performed using GEPIA2 [27], a web server that contains RNA sequencing data from TCGA and GTEx projects [72], [73]. The ANOVA method was employed to obtain differential expression data for all genes under study, depicting the mRNA expression in tumor (T) relative to corresponding normal (N) tissues – HCC (T:n=369; N:n=160), BRCA (T:n=1085; N:n=291), LUAD (T:n=483; N:n=347). Next, subcellular localization of differentially expressed CSPs was retrieved from online protein databases, such as UniProt [28] and Gene Ontology [29]. Focusing on MPZL1 as a CSP target, the correlation between its CNV status and mRNA expression levels in the same datasets was explored via cBioPortal considering only samples with both data types available (HCC, n=360; BRCA, n=960; LUAD, n=230). Additionally, the CNV status of MPZL1 in human cancer cell lines was retrieved from the cancer cell line encyclopedia [74].

### Cell culture of human established cell lines

All cell lines were incubated at 37°C with 5% CO_2_ and maintained in sterile conditions. Depending on the specific cell line, high glucose Dulbecco′s Modified Eagle′s Medium (DMEM, Sigma-Aldrich) or Roswell Park Memorial Institute Medium (RPMI 1640, Gibco^TM^) were used, in every case supplemented with 10% FCS (Gibco^TM^) and 1% penicillin/streptomycin (10,000 U/mL penicillin and 10 mg/mL streptomycin, Sigma-Aldrich). All human liver and breast cancer cell lines, as well as U-87 MG and CFPAC-1, were cultured in DMEM medium, while all human lung cancer cell lines, as well as NALM6 and Molm13, were cultured in RPMI medium.

### Genetic modification of cancer cell lines

For CRISPR/Cas9-mediated knockout of MPZL1, sgRNAs were designed using the CHOPCHOP web tool [75], and subsequently cloned into the pLentiCRISPR v2 vector [76], as previously described [77]. For lentiviral overexpression of MPZL1, NEBuilder^®^ HiFi DNA Assembly was used to introduce a MPZL1 PCR product, generated from pENTR223-MPZL1 (obtained from DKFZ core facility), into the pLenti6.2/V5-DEST vector. Lentivirus production, transduction of cancer cells, and antibiotic-based selection were performed as previously described [77], generating isogenic cancer cell lines either lacking or overexpressing MPZL1 protein. Finally, the efficiency of genetic modification was assessed via immunoblotting.

- sgMPZL1.1_top guide (5’ → 3’): CACCGGGGGGCCGACACTACTGT
- sgMPZL1.1_bottom guide (5’ → 3’): AAACACACAGTAGTGTCGGCCCC
- sgMPZL1.2_top guide (5’ → 3’): CACCGGCACCGACCACAGCCAGC
- sgMPZL1.2_bottom guide (5’ → 3’): AAACGCGCTGGCTGTGGTCGGTG

### Immunoblotting

Harvested cells were lysed in lysis buffer (Cell Signaling Technology), supplemented with protease (cOmplete^TM^ Mini, Roche) and phosphatase inhibitors, by incubation on ice for 30 minutes, followed by centrifugation at 13,000 rpm to collect protein lysates. Quantification of protein was done using the Bradford reagent (Bio-Rad). Samples with equal concentration were the prepared with Laemmli buffer [77] and denatured by boiling at 95°C for 5 minutes.

20 μg protein samples were separated via SDS-PAGE using manually casted polyacrylamide gels and EZ-Run™ protein ladder (Fisher BioReagents™) as indicator of the molecular weight. Separated proteins were transferred to ROTI^®^PVDF membranes (Carl Roth) via Western Blot. Membranes were then blocked for 1 hour in 5% Milk (Carl Roth), followed by incubation with the primary antibody overnight at 4°C (Table 1). The day after, membranes were washed with TBS/T buffer and incubated with the corresponding HRP-conjugated secondary antibody for 1 hour at RT, followed by additional washing and chemiluminescent signal detection using the Clarity™ Western ECL Substrate (Bio-Rad) and ChemiDoc^TM^ Gel Imaging System (Bio-Rad). Quantification of protein bands was done via densitometry-based analysis with ImageJ, where the relative intensity of MPZL1 was normalized to that of Vinculin, used as loading control.

**Table 1.**
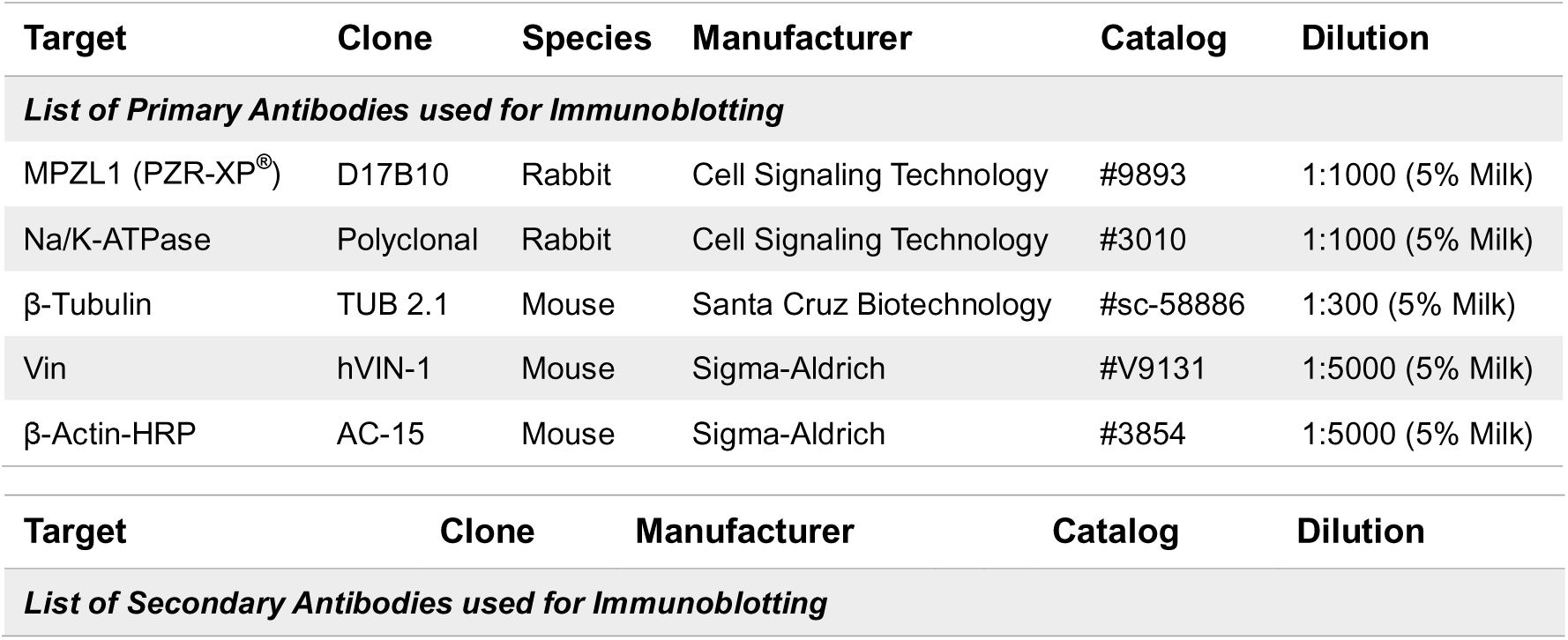
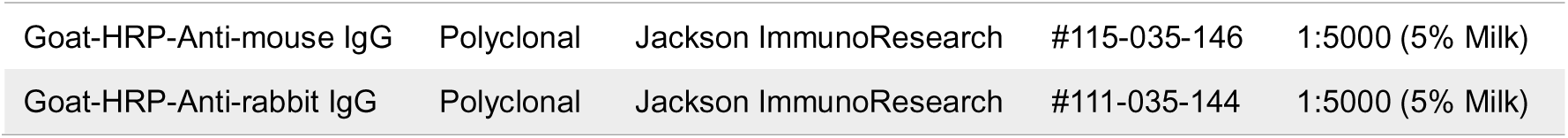
Antibodies used for immunoblotting.

### Deglycosylation assay

Deglycosylation of protein lysates was achieved by treatment with PNGase F enzyme. First, denaturation reactions were prepared by mixing 20 µg of lysate and 1 µL of Glycoprotein Denaturing Buffer (NEB) in a final volume of 10 µL of H_2_O. Samples were then boiled at 100°C for 10 minutes. Second, deglycosylation reactions were set up by adding 2 µL of GlycoBuffer (NEB), 2 µL of NP-40 (NEB), 6 µL of H_2_O, and 1 µL of PNGase F (NEB), followed by incubation at 37°C for 1 hour. Non-deglycosylated samples were subjected to the same denaturation protocol as internal experimental control. Samples were further analyzed via immunoblotting.

### Proliferation assays

For the clonogenic assay, 1×10^3^ cells were plated in triplicates into 6-well plates and allowed to grow during two weeks, with regular medium exchange. The cell colonies were then fixed and stained with a 0.05% (w/v) crystal violet solution containing 1% formaldehyde. Plates were scanned to generate digital images using the Perfection V850 Pro scanner (Epson). For the short-term proliferation assay, 1×10^3^ cells were seeded in triplicates into independent 96-well plates corresponding to the different time points (0, 24, 48, 72, and 96 hours). The readout was performed every 24 hours using CellTiter-Blue^®^ (Promega), following the manufacturer’s instructions. After incubation during 4 hours at 37°C, fluorescence intensity (560/590 nm) was measured using the EnSpire^®^ Multimode Plate Reader (PerkinElmer). *Time 0* was defined as the day after plating the cells, being the results normalized to this time point.

### Cell fractionation assay

To investigate the subcellular localization of MPZL1, the Minute^TM^ Plasma Membrane Protein Isolation and Cell Fractionation Kit (Invent Biotechnologies) was used. Cells grown in 15 cm dishes were harvested and processed according to the manufacturer’s protocol. After multiple rounds of centrifugation and incubation in the buffers provided, the different protein fractions were obtained: cytosol, organelle membrane, and plasma membrane. Additionally, *“whole cell lysates”* from the same cell lines were prepared as experimental controls. Samples were analyzed via immunoblotting, using tubulin and Na/K-ATPase as markers for cytosolic and plasma membrane fractions, respectively.

### MPZL1 CSP target in a human patient cohort

Thirteen samples of patient livers were obtained upon tissue surgical resection [78], [79]. Copy number variation status of MPZL1 gene was assessed via comparative genomic hybridization (CHG). Next, MPZL1 mRNA expression was evaluated with microarray analysis, while MPZL1 protein levels were assessed through immunoblotting.

### Immunohistochemistry

IHC was performed as previously described [77]. Briefly, deparaffinization of tissues was achieved by incubating the slides in xylene, followed by rehydration using a descending series of alcohol and a washing step with water. For antigen retrieval, slides were boiled for 8 minutes in a pressure cooker using a sodium citrate buffer (10 mM trisodium citrate dihydrate, 0.5%(v/v) TWEEN 20, pH 6.0), followed by rinsing in water for 5 minutes to cool down. Next, endogenous HRP was blocked by a 10-minute incubation in 3% hydrogen peroxide diluted in H_2_O. The slides were then rinsed in water for another minute, and washed twice for 2 minutes with PBS. Tissue sections were blocked for 1 hour at room temperature in 5% BSA (Carl Roth) containing 0.5% Triton X-100, followed by overnight incubation at 4°C with the corresponding primary antibody diluted in blocking buffer (Table 2). Then, the slides were washed three times with PBS containing 0.05% Triton X-100 for 5 minutes, and incubated with the corresponding ImmPRESS^®^ HRP Horse IgG Polymer Detection Kit, Peroxidase (Anti-Rabbit or Anti-Mouse) (VectorLabs) during 30 minutes at RT, again followed by three 5-minute washing steps with PBS/Triton X-100. The antibody signal was detected with ImmPACT^®^ DAB/HRP substrate (VectorLabs), according to the manufacturer’s instructions, with the reaction time adjusted individually for each antibody. Counterstaining with hematoxylin solution (Carl Roth) was then carried out for 1 or 2 minutes to obtain improved staining contrast. The slides were rinsed with running water and dehydrated using an ascending alcohol series, finishing with xylene. When dried, the coverslips were mounted on the slides using Micromount^®^ mounting media (Leica). Histological slides were scanned with the Hamamatsu NanoZoomer Digital Pathology (NDP) system, and visualized and analyzed using QuPath [80] and FiJi ImageJ [81] software.

**Table 2.**
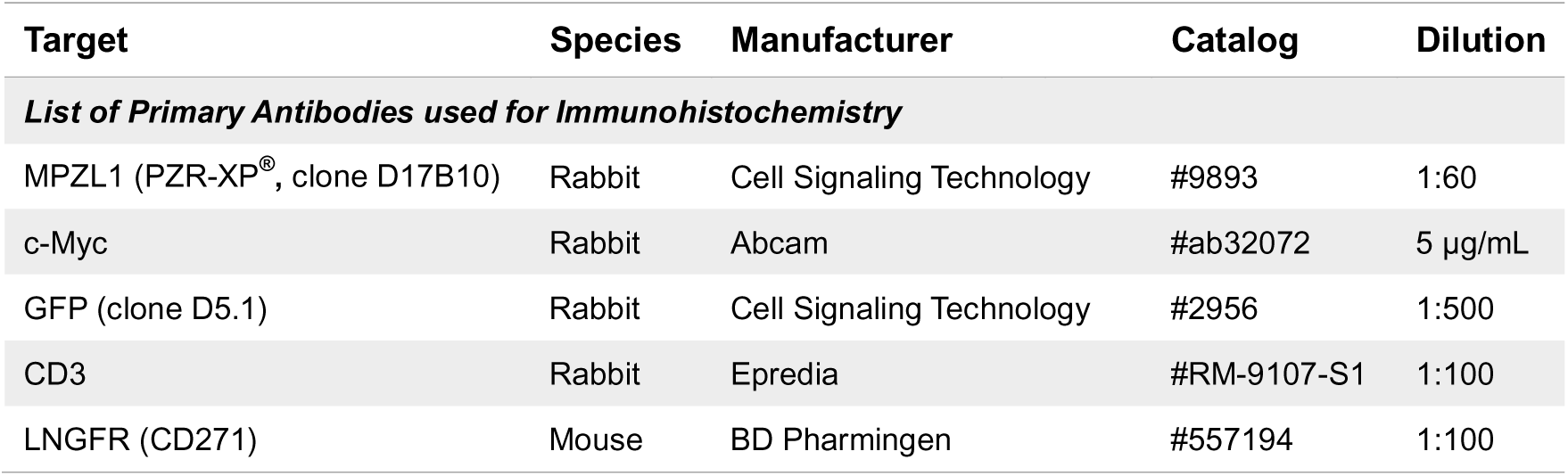
Antibodies used for immunohistochemistry.

### Tissue microarray (TMA) analysis

The expression and subcellular localization of MPZL1 in primary human tissue was analyzed by immunohistochemistry using TMAs, comprising cancer-specific and non-cancer tissue punches (Table 3). The histochemical staining was quantified by an experienced pathologist using a scoring system to evaluate MPZL1 expression (MPZL1-score) on the surface of epithelial cells, where 0 indicates negative staining, 1 and 2 represent intermediate levels, and 3 corresponds to complete plasma membrane staining in more than 10% of the cells (Figure 2E). Institutional Ethical Review Board approval was obtained at local Ethical Committees of the Medical Universities of Heidelberg (S205-06), in compliance with the Helsinki Declaration. Written informed consent was obtained from all individuals.

**Table 3.**
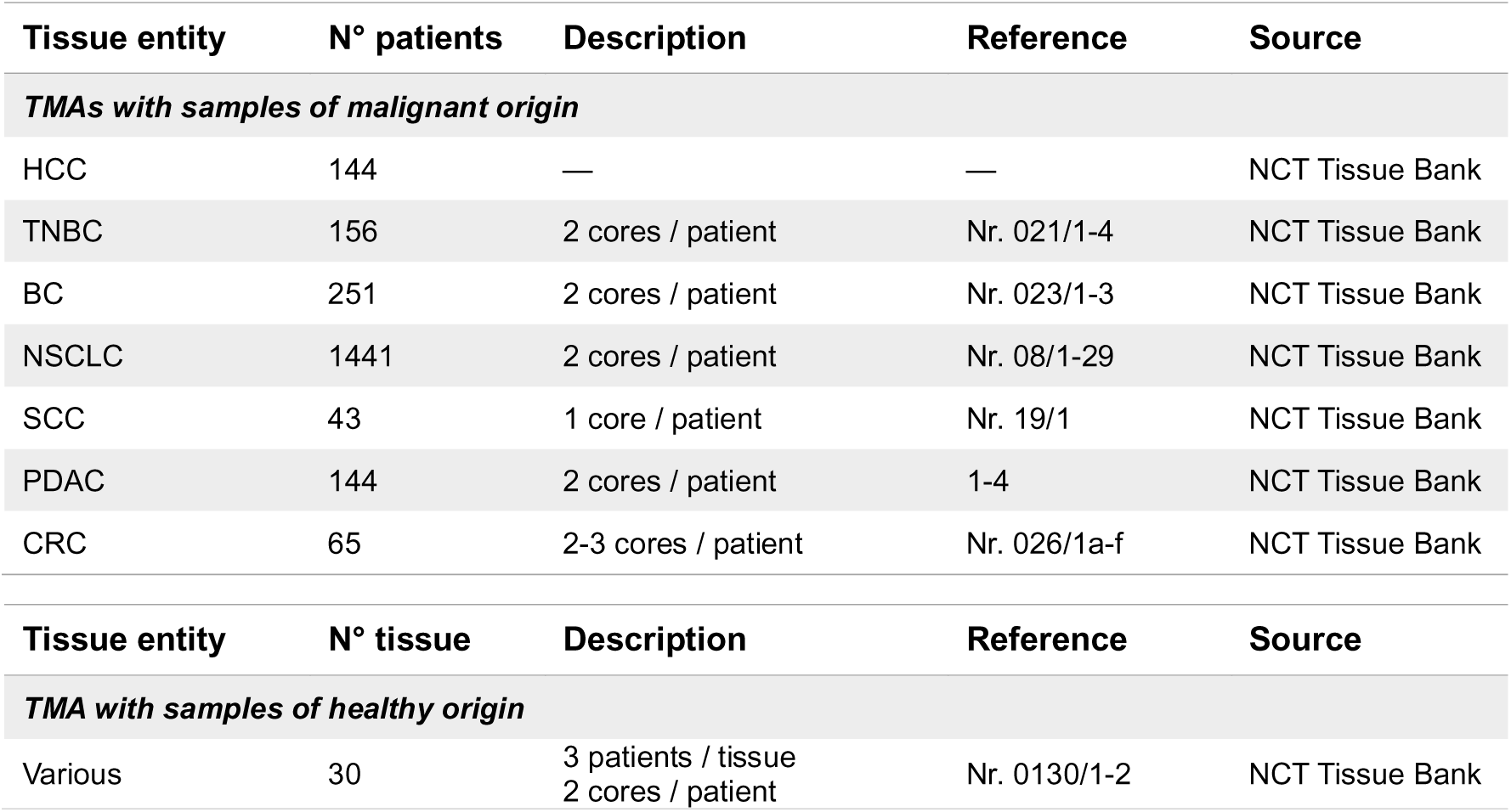
Human TMAs used for MPZL1 IHC.

| Tissue entity | N° patients | Description | Reference | Source |
| --- | --- | --- | --- | --- |
| *TMAs with samples of malignant origin* |  |  |  |  |
| HCC | 144 | — | — | NCT Tissue Bank |
| TNBC | 156 | 2 cores / patient | Nr. 021/1-4 | NCT Tissue Bank |
| BC | 251 | 2 cores / patient | Nr. 023/1-3 | NCT Tissue Bank |
| NSCLC | 1441 | 2 cores / patient | Nr. 08/1-29 | NCT Tissue Bank |
| SCC | 43 | 1 core / patient | Nr. 19/1 | NCT Tissue Bank |
| PDAC | 144 | 2 cores / patient | 1-4 | NCT Tissue Bank |
| CRC | 65 | 2-3 cores / patient | Nr. 026/1a-f | NCT Tissue Bank |
| Tissue entity | N° tissue | Description | Reference | Source |
| *TMA with samples of healthy origin* |  |  |  |  |
| Various | 30 | 3 patients / tissue<br>2 cores / patient | Nr. 0130/1-2 | NCT Tissue Bank |

### Generation of an antibody targeting MPZL1

The hybridoma technology was used to generate antibodies against the extracellular domain of MPLZ1 protein. Briefly, mice were immunized with the extracellular fragment of human MPZL1 protein (aa 36-165). When an animal developed strong titers, its B lymphocytes were isolated and fused with immortal myeloma cells to generate hybridoma clones capable of producing unique monoclonal antibodies (mAbs), and the clones were evaluated via FACS. The identification of a positive clone was followed by two rounds of subcloning, and the most specific subclone against MPZL1 was selected. Subsequently, the RNA from this hybridoma was sequenced to identify the variable heavy (V_H_) and *kappa* light (V_L_) chains (Table 4). These sequences were cloned in frame into two independent human backbone vectors coding for the constant heavy (C_H_) and *kappa* light (C_L_) chains [40], respectively. Finally, PEI-mediated transfection of HEK293 cells with these two plasmids led to the production of MPZL1-ChAb, composed of a human constant domain and a mouse variable region.

**Table 4.**
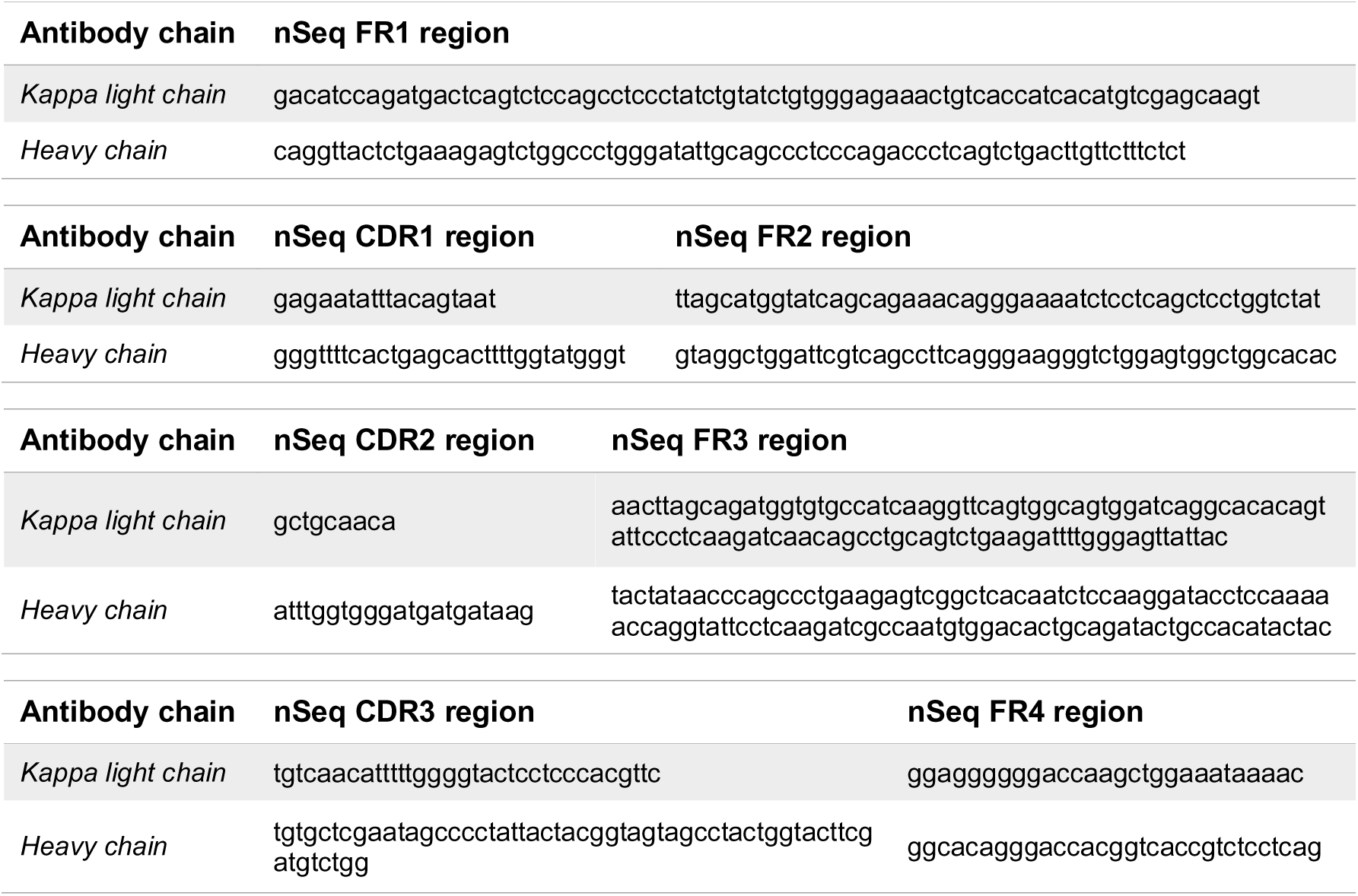
Sequence of the variable regions of the MPZL1 hybridoma. Antibody chain nSeq FR1 region.

### Cell viability assay

To evaluate cellular susceptibility to MPZL1-ChAb, 1×10^3^ cells per well were seeded in a 96-well plate and allowed to attach for 24 hours. Consecutively, treatments with increasing concentrations of MPZL1-ChAb (0,05 – 100 μg/mL) were applied in triplicates, with DMEM medium as negative control. The readout was performed after 72 hours using CellTiter-Blue^®^ (Promega), following the manufacturer’s instructions. Fluorescence intensity (560/590 nm) was measured with the EnSpire^®^ Multimode Plate Reader (PerkinElmer).

### Cell surface staining and FACS analysis

Extracellular staining of MPZL1 in isogenic cancer cell lines was performed to evaluate the specificity of MPZL1-ChAb. Briefly, adherent cells were washed with PBS and incubated in Versene-EDTA (Lonza) at RT for 10 to 15 minutes prior to harvesting using FACS buffer (3% FCS in PBS). 3×10^5^ cells per condition were transferred to previously annotated wells in a V-bottom 96-well plate. Upon washing, cells were stained with 100 µg/mL MPZL1-ChAb for 45 minutes at 4°C. Next, cells were washed and stained with a secondary antibody (AF488^®^ or APC anti-human IgG Fc, BioLegend) during 45 minutes at 4°C in the dark. After a final washing step, cells were resuspended in 200 µL FACS buffer containing 1:1000 SYTOX^®^ Blue (Invitrogen), and transferred to cell strainer snap cap FACS tubes (Corning) for FACS analysis using a BD LSRFortessa flow cytometer (BD Biosciences). As negative controls for specificity, all the experimental samples were stained in parallel with human IgG1 isotype (BioLengend) at the same concentration as the primary antibody. When CD19 staining was included, the samples underwent a third incubation round with PE anti-human CD19 antibody (BioLegend) during 20 min in the dark. The same protocol was followed to evaluate MPZL1 expression in cell lines from different cancer types, as well as in CAR-T cells. Data analysis was in all cases carried out with FlowJo v10.

### FFPE tissue blocks and matched DTCs

Cryopreserved dissociated tumor cells (DTCs) and formalin-fixed paraffin-embedded (FFPE) tissue blocks from human breast tumors of five independent patients (Discovery Life Sciences) were used to investigate MPZL1 expression on the surface of cancer cells within the tumor architecture. Anonymous patients were named as: 8843, 4130, 8921, 2457, 8991. MPLZ1 expression on tissue slides obtained from the FFPE blocks was evaluated via IHC with MPZL1 commercial antibody. DTCs were thawed and stained with fluorochrome-conjugated MPZL1-ChAb to measure antibody binding capacity on the cell surface via FACS analysis.

### Measurement of antibody binding capacity

To quantitatively measure the antibody binding capacity (ABC) in DTCs, both MPZL1-ChAb and human IgG isotype control (Invitrogen) were covalently labeled with the AF488^®^ fluorochrome, using the AlexaFluor^TM^ 488 protein labeling kit (Invitrogen), according to the manufacturer’s instructions. Following labeling and purification, the sample absorbance was determined utilizing a Nanodrop ND-1000 Spectrophotometer (Thermo Fisher Scientific), with wavelengths of 280 nm for the protein and 494 nm for the dye. The protein concentration and the degree of labeling (DOL), representing the moles of dye bound per mole of protein, where calculated following protocol provided in the kit. For cell surface staining, 1×10^5^ DTCs per condition were used, with JIMT-1 and Molm13 cells included as positive and negative controls, respectively. All samples were first incubated with 5 µg/mL of Human BD Fc Block^TM^ (BD Pharmingen^TM^) for 10 minutes at RT to minimize nonspecific antibody binding to Fc receptors. Directly afterwards, cells were incubated with 5 µg/mL of labeled antibody for 1 hour at 4°C in the dark. After two washing steps with PBS, cells were resuspended in 200 µL FACS buffer containing 1:1000 of SYTOX^®^ Blue (Invitrogen). Samples were acquired in a BD LSRFortessa flow cytometer (BD Biosciences). A calibration curve was built to correlate the instrument’s acquisition channels to standardized fluorescence intensity units known as MESF (molecules of equivalent soluble fluorochrome), employing the Quantum™ Alexa Fluor^®^ 488 MESF kit (Bangs Laboratories), according to the manufacturer’s instructions. This calibration enabled the interpolation of MESF values from the median channel values obtained from the acquired samples. Finally, the ABC in each sample was calculated dividing the MESF value by the DOL value previously determined. Additionally, the ABC for the IgG isotype control was subtracted from that of the MPZL1-ChAb in each sample to account only for specific target binding. This method allowed to determine the binding efficiency of MPZL1-ChAb to patient cancer samples with various levels of MPZL1 expression on the cell surface of cancer cells.

### Generation of MPZL1-specific CAR constructs

The scFv generated is composed of the variable regions of the heavy and light chains of MPZL1-ChAb (Table 4), connected by a short linker peptide. Two distinct approaches were followed, resulting in V_L_+V_H_ and V_H_+V_L_ configurations, with two constructs designed for each combination. The MPZL1-scFv is preceded by a leader sequence and followed by a hinge-transmembrane region, along with the CD28 and CD3ζ domains. The construct also includes the expression of a reporter gene (LNGFR). Briefly, these components were incorporated into a SFGγ retroviral vector, which was used for the stable transfection of gpg29 (H29) fibroblasts. The generated retroviral supernatants were employed to establish stable retrovirus-producing cell lines, being the resulting supernatants utilized for the transduction of T cells [82].

### Isolation and expansion of human T cells

Buffy coats from anonymous healthy human donors were purchased from either the ZKT Tübingen or the IKTZ Heidelberg (in German, *Zentrum für Klinische Transfusionsmedizin,* and *Institut für Klinische Transfusionsmedizin und Zelltherapie*, respectively). Peripheral blood mononuclear cells (PBMCs) were separated with Lymphocyte Separation Medium (Corning) via density gradient centrifugation. Isolated PBMCs were cryopreserved in FCS containing 10% DMSO. T cells were purified from thawed PBMCs by negative selection using magnetic-activated cell sorting (MACS) with a Pan T Cell Isolation Kit (Miltenyi Biotec), following the manufacturer’s instructions. Isolated T cells were stimulated with CD3/CD28 human T cell Activator Dynabeads (Gibco) in a 1:1 ratio, and plated at 1×10^6^ cells per mL in RPMI medium supplemented with FCS and penicillin/streptomycin, and 5 ng/mL of interleukin-7 (IL7) and interleukin-15 (IL15) cytokines (Miltenyi Biotec). Specific stimulation continued for 72 hours before proceeding to T cell genetic modification. Cells were in all cases maintained in sterile conditions and incubated at 37°C with 5% CO2.

### Genetic modification of T cells

After 72 hours in culture, the Dynabeads were magnetically removed from the activated T cells. Subsequent T cell transduction was performed utilizing recombinant human fibronectin fragment (RetroNectin^®^, Takara Bio), which aids in the co-localization of target cells and viral particles when subjected to spinoculation. Briefly, non-tissue culture treated multi-well plates were coated with 15 µg/mL RetroNectin^®^ diluted in PBS for 2 hours at RT. The coating solution was then removed and the following transduction components were added: (1) CAR-coding retroviral supernatant, (2) T cells at a concentration of at 1M T cells/mL supplemented with 5 ng/mL of IL7 and IL15 cytokines, and (3) RPMI medium to reach the same final volume across all conditions. Spinoculation took place for 1 hour at 1,200 rpm and 37°C. Upon transduction, the cytokine-supplemented medium was replenished every second day.

### CAR expression, proliferation and viability of unstimulated CAR-T cells

CAR-T cells are defined as *unstimulated* when they have not encountered their specific antigen, and are maintained in culture in the presence of IL7 and IL15 cytokines. Expression of the CAR construct on the surface of T cells was determined via flow cytometry starting from day four after transduction. T cells were stained with both AF647^®^ goat anti-mouse IgG F(ab’)2 fragment specific (AffiniPure) and PE mouse anti-human CD271 (LNGFR, BD Biosciences). Cell acquisition was performed on a Cytek^®^ Aurora (Cytek Biosciences) or a LSRFortessa (BD Biosciences) flow cytometer, and subsequent data analysis was carried out with FlowJo v10. For proliferation and viability assays, T cells were counted using an automated cell counter (Countess™, Invitrogen) at multiple time points after transduction, and viability percentages were automatically calculated based on trypan blue exclusion.

### Bioluminescence-based cytotoxicity assays

To generate target cells, isogenic cancer cell lines were stably transduced to express firefly luciferase (FFLuc) linked to GFP protein. The GFP+ population was sorted using a BD FACSAria™ Fusion cell sorter (BD Biosciences) to have a pure population of Luc-expressing cells. For the cytotoxic assays, 2×10^4^ adherent target cells per well were seeded in black-walled transparent-bottom 96-well plates and incubated at 37°C and 5% CO_2_ during 24 hours to allow for attachment. After that, medium was removed and the co-culture with CAR-T cells was established in triplicates at the indicated range of effector-to-target cell ratios (E:T ratio), in a final volume of 100 µL/well. In every case, the required amount of T cells was calculated based on their percentage of CAR-positive T cells, adjusting the total number of effector cells across treatments via compensation with UT T cells. Target cells alone and 0.2% Triton X-100 were utilized for the minimal and maximal lysis controls (RLU, relative light units), respectively. In the case of target suspension cells, no pre-seeding was required. Following the co-culture of cells during 4 or 18 hours (short- and long-term killing assays, respectively), 100 µL of D-luciferin (ZellBio, GoldBio) diluted 1:10 in PBS were added per well. Bioluminescence was measured using a microplate reader (Infinite^®^ 200 PRO, Tecan), and cell lysis was determined as (1−(RLU_sample_)/(RLU_max_))×100.

### Phenotypic characterization

For phenotypic analysis, flow cytometry was performed after staining MPZL1 CAR-T cells with the following mouse anti-human fluorochrome-conjugated antibodies: BUV395 CD4, BV650 CD45RA, and PE CD271 (LNGFR) (BD Biosciences); PE/Cyanine7 CD8a, and PerCP-eFluor 710-LAG3 (Invitrogen); and BV510 CD25, BV711 CD69, APC/Cyanine7 PD-1, and BV421 CD62L (BioLegend), along with 7-AAD cell viability solution (BD Biosciences). Besides, compensation controls using UltraComp eBeads^TM^ (Invitrogen) and fluorescence minus one (FMO) controls were included. Cell acquisition was done on a Cytek^®^ Aurora flow cytometer (Cytek Biosciences), and subsequent data analysis was carried out with FlowJo v10.

### Antigen-dependent proliferation and intracellular cytokine secretion assay

Antigen stimulation assays consisted in the co-culture of MPZL1 CAR-T cells with either MPZL1^high^ JIMT-1 breast cancer cells or the respective *MPZL1* knockout cells. For the antigen-dependent proliferation assay, JIMT-1 target cells were irradiated with a dose of 30 grays during 4 minutes to inhibit their proliferative capacity prior to the co-culture. Next, 3×10^5^ irradiated cells per well were seeded in 24-well plates, and incubated during 24 hours to allow for attachment. DMEM was removed after one day, and the co-culture was established in triplicate wells by adding 1×10^6^ MPZL1 CAR+ T cells without cytokines in 500 µL of RPMI per well, with fresh RPMI added every other day. T cell counts were performed every six days with a cell counter (Countess™, Invitrogen). Subsequently, the co-culture was reestablished by adding 1×10^6^ prestimulated CAR-T cells (after a 6-day stimulation period) to freshly irradiated target cells. Restimulation was conducted over three consecutive 6-day periods, with the cumulative number of CAR+ T cells calculated as the product of the current CAR-T cell count and the accumulated expansion factor, which incorporates the growth rates of previous stimulations. For the intracellular cytokine secretion assay, a 16-hour co-culture of 5×10^5^ MPZL1 CAR-T cells and 5×10^5^ respective JIMT-1 target cells, seeded one day in advance, was established in 24-well plates. Treatment with PMA/Ionomycin was utilized as a positive control to induce cytokine secretion. During the last 4 hours, a cocktail of Brefeldin A and Monensin (protein transport inhibitor cocktail, Invitrogen) was added to the co-cultures to retain the cytokines within the cells. Next, T cells were washed, stained for cell surface markers (phenotypic analysis), fixed and permeabilized with a fixation and permeabilization buffer set (Invitrogen), according to the manufacturer’s instructions. Intracellular cytokine staining was performed with the following antibodies: APC Mouse Anti-Human/Mouse GZMB and PerCP Rat Anti-Human IL-2 (BioLegend), and BUV737 Mouse Anti-Human IFNγ and BV650 Mouse Anti-Human TNFα (BD Biosciences), along with ViaDye™ Red Fixable Viability Dye (Cytek Biosciences). Both compensation and FMO controls were included. Cell acquisition was done on a Cytek^®^ Aurora flow cytometer (Cytek Biosciences), and subsequent data analysis was carried out with FlowJo v10.

### Mouse experiments

MPZL1 CAR-T cells used for animal treatments were always administered directly after primary *ex vivo* expansion. Tumor formation was induced in all cases in 7-to 8-week-old mice, and all animal experiments were conducted in compliance with the regional regulations and with the approval of the regional board (Regierungspräsidium Karlsruhe, Germany), under the permit number G-70/20.

### Human cell line-derived mouse xenograft model

Tumors of human origin were established in immunodeficient NSG mice (Janvier). To this end, 5×10^6^ genetically manipulated cells in 100 μL of PBS were subcutaneously injected into each flank of the mouse under isoflurane anesthesia. The xenograft tumors were regularly measured with a caliper, and tumor volume was calculated as (length×(width)^2^)/2. When the average tumor volume reached a specific size (120 mm^3^ for breast and 360 mm^3^ for HCC xenografts), a single dose of 1×10^7^ MPZL1 CAR-T cells was intravenously administered to all the animals in the experiment on the same day (time 0). Tumor size was monitored over time, and the mice were euthanized by cervical dislocation when any of their tumors reached a maximum volume of 1,000-1,500 mm^3^ or, alternatively, if ulcerations appeared. The tumors were resected, photographed, weighed, and finally fixed in 4% paraformaldehyde (PFA) for further processing in IHC applications.

### HTVI-induced autochthonous HCC mouse model

Tumors were generated directly in the livers of NSG mice by hydrodynamic tail vein injection (HTVI), where 2 mL of NaCl solution (approximately 10% of the mouse body weight) containing naked plasmid DNA was rapidly injected (within 5 to 7 seconds) into the tail vein, enabling efficient transfection of hepatocytes in vivo [83]. Tumor formation was induced by overexpression of c-MYC, coupled with either human MPZL1 cDNA or GFP, employing the Sleeping Beauty DNA transposon system [84], and CRISPR/Cas9-mediated KO of p53, with the respective plasmids generated as previously described [77]. The groups were respectively named MycOE;p53-/-;MPZL1 and MycOE;p53-/-;GFP. For the intervention experiment, a single dose of 1×10^7^ MPZL1 CAR-T cells was intravenously administered to all the animals 15 days after HTVI, and formation of tumors was evaluated by regular palpation of the mice bellies. When tumors were detected, the mice were euthanized by cervical dislocation. The livers were resected, acquiring bright-field and GFP pictures using a stereomicroscope (MZ10F, Leica), and fixed in 4% PFA to be processed for IHC studies. In the short-term experiment, a single dose of 1×10^7^ MPZL1 CAR-T cells was intravenously administered one month after HTVI. One week after CAR-T treatment, all animals were sacrificed and the livers underwent the same processing.

### Statistics

If not indicated otherwise, statistical analyses were performed using GraphPad Prism 10 (Version 10.0.0).

## Supplemental Figures

**Suppl. Figure S1.**
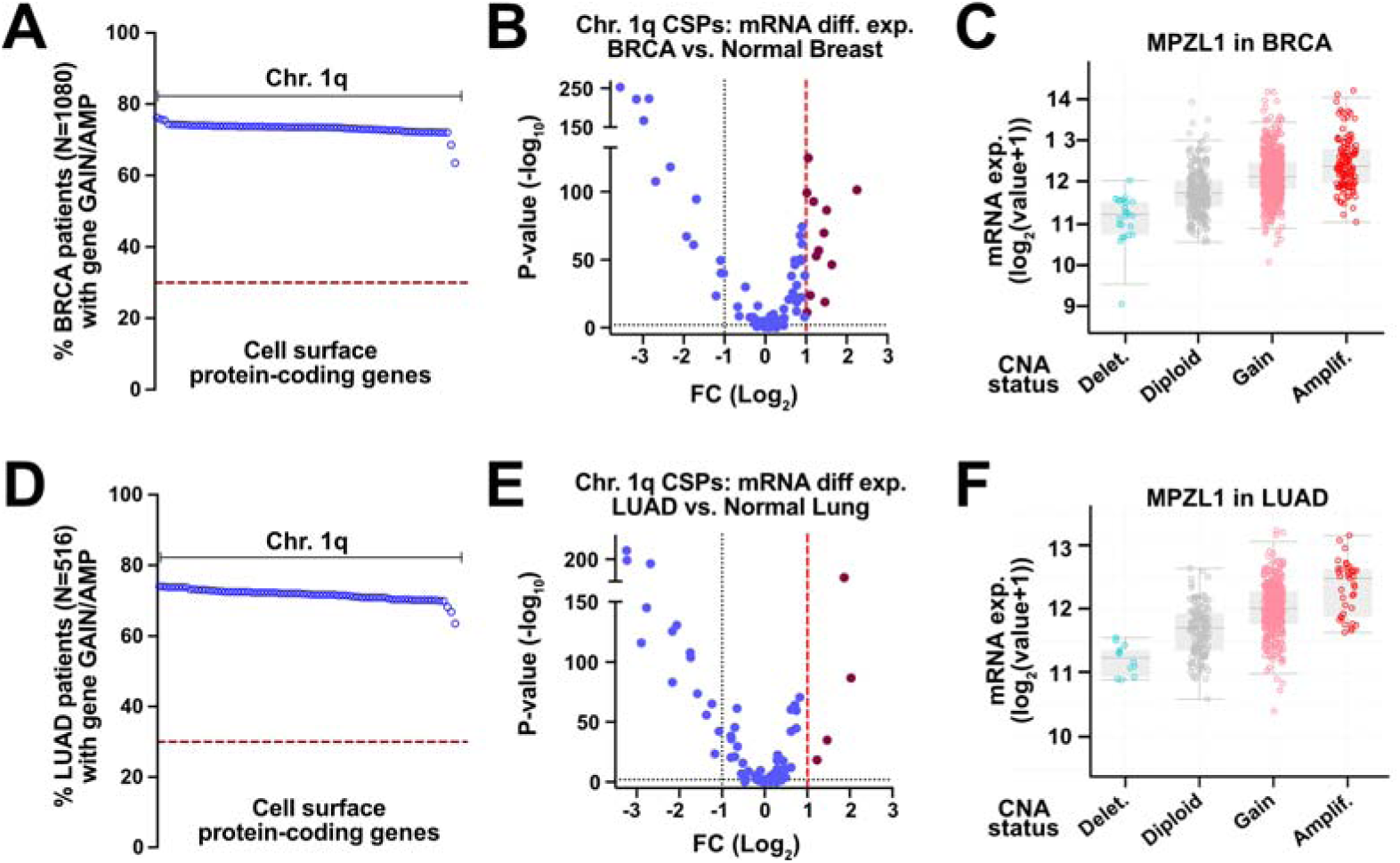
CSP analyses in cancer entities with recurrent chr. 1q amplifications. (A) – (c) Breast cancer. **(A)** Percentage of BRCA patients (n=1080) who exhibit a gain (score 1) or amplification (score 2) of each of the hits defined in Figure 1B. **(B)** Volcano plot depicts mRNA differential expression data of chr. 1q frequently amplified CSP-coding genes in breast tumors compared to normal breast tissues (Tumor:n=1085; Normal:n=291). **(C)** Boxplots depict the correlation of MPZL1 CNV status and its mRNA expression levels in the BRCA TCGA dataset (n=960). **(D) – (F) Lung cancer. (D)** Percentage of LUAD patients (n=516) who exhibit a gain (score 1) or amplification (score 2) of each of the hits. **(E)** Volcano plot shows mRNA differential expression data of chr. 1q frequently amplified CSP-coding genes in lung tumors compared to normal lung tissues (Tumor:n=483; Normal:n=347). **(F)** Boxplots depict the correlation of MPZL1 CNV status and its mRNA expression levels in the LUAD TCGA dataset (n=230). Analyses in (A), (C), (D) and (E) are performed with cBioPortal. CNV status is defined by the GISTIC2.0 algorithm, which assigns a specific score to each gene in each patient: -1 (shallow deletion), 0 (diploid), 1 (gain), and 2 (amplification). mRNA data is quantified using the expectation-maximization algorithm and expressed as log_2_(value+1). Analyses in (B) and (E) are done using the GEPIA2 online resource and the ANOVA method. Data is expressed as log_2_(fold-change) with cut-off set at log_2_(FC)>1.

**Suppl. Figure S2.**
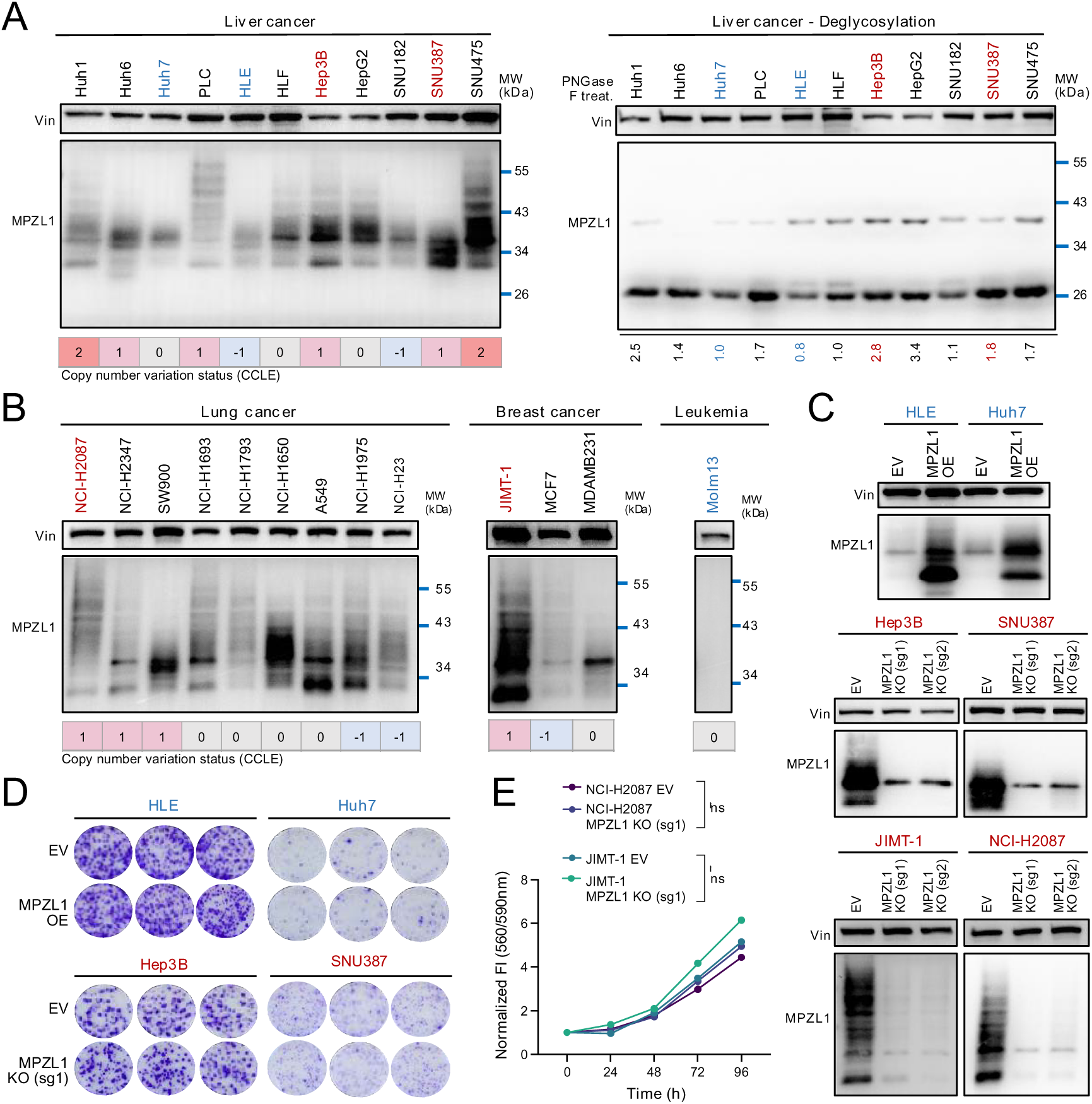
MPZL1 expression in established human cell lines from solid cancer entities. **(A) Left panel**: immunoblotting analysis of whole cell lysates (WCLs) from human liver cancer cell lines explores MPZL1 expression levels. CNV data for these cell lines is retrieved from the Cancer Cell Line Encyclopedia (CCLE) using cBioPortal. CNV status is defined by the GISTIC2.0 algorithm and depicted as color-coded numbers: red (amplification: score 2), pink (gain: score 1), gray (diploid: score 0) and light blue (shallow deletion: score -1). **Right panel**: WCLs are subjected to deglycosylation by treatment with PNGase F, followed by MPZL1 protein expression analysis via immunoblotting. Vinculin is used as loading control. Numbers below the blot represent densitometry-based analysis of MPZL1 protein band intensities normalized to Vinculin for each sample (quantification done with ImageJ). **(B)** Immunoblotting analysis of MPZL1 expression in WCLs from several human breast and lung cancer cell lines. CNV status is defined as in (A). Leukemia Molm13 cell line is used as negative control. Red and blue colours identify, respectively, the cell lines selected as MPZL1^high^ and MPZL1^low^. **(C) – (E) Generation and characterization of isogenic cell lines. (C)** Immunoblotting analysis of MPLZ1 expression. Vinculin is used as loading control. MPZL1^low^ cancer cell lines are stably transduced with either pLenti6.2/DEST-V5 empty vector (EV) or same vector carrying human *MPZL1* cDNA for overexpression (OE). MPZL1^high^ cancer cell lines are stably transduced with either LentiCRISPR V2 empty vector (EV) or same vector harboring a sgRNA targeting *MPZL1* gene to knock it out – two different guides are used (named as sg1 and sg2). **(D)** Colony formation assays of HCC isogenic cell lines. Briefly, 1×10^3^ cells/well are plated in triplicates into 6-well plates and allowed to grow for two weeks before fixation and staining with crystal violet solution. Digital images are generated using the Perfection V850 Pro scanner (Epson). **(E)** Short-term cell proliferation assays of isogenic cell lines. Briefly, 1×10^3^ cells/well are seeded in triplicates into independent 96-well plates for the different time points. Experimental readout is made through CellTiter-Blue^®^ every 24 hours, using a multiplate reader to measure fluorescence intensity (560/590 nm). Paired t-test performed for statistical analysis (ns: nonsignificant).

**Suppl. Figure S3.**
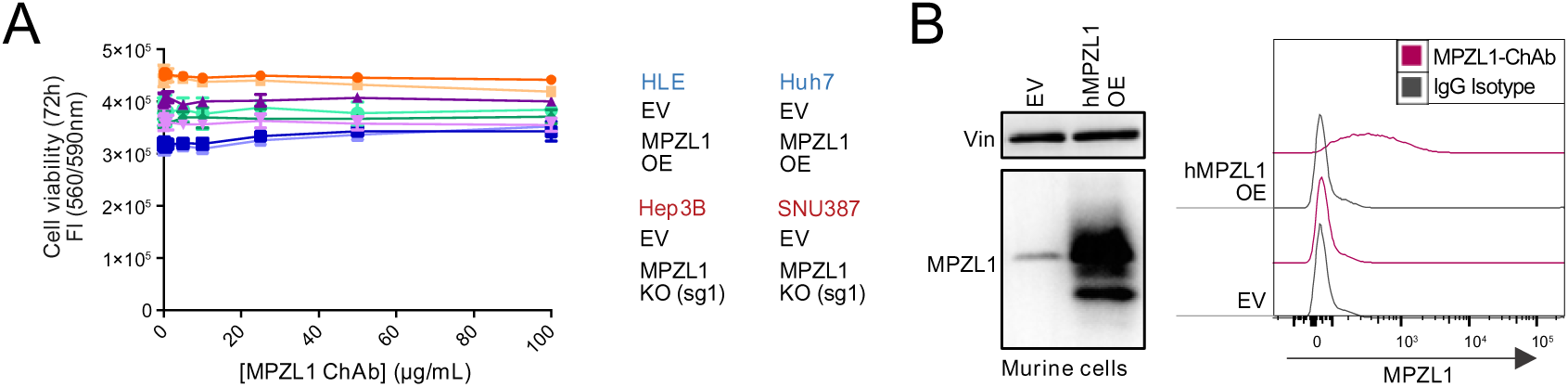
Further characterization of MPZL1-ChAb. **(A)** Evaluation of cellular susceptibility to MPZL1-ChAb using isogenic liver cancer cell lines. Briefly, 1×10^3^ cells/well are seeded in a 96-well plate and treated with increasing concentrations of MPZL1-ChAb (0.05 – 100 µg/mL) in triplicates, with normal DMEM used as negative control. Experimental readout is performed via CellTiter-Blue^®^ after 72 hours, measuring fluorescence intensity (560/590 nm) in a microplate reader. **(B)** Evaluation of MPZL1-ChAb binding to murine cells. Liver cancer cells derived from HTVI-induced genetically defined tumors with the KrasG12D;p53-/-genotype are used. Murine cells are transduced with either pLenti6.2/DEST-V5 empty vector (EV) or same vector carrying human *MPZL1* cDNA for overexpression (hMPZL1 OE). **Left panel**: MPZL1 protein expression is analyzed by immunoblotting using the commercially available antibody, which specifically binds human MPZL1, but not the murine ortholog. Additionally, endogenous MPZL1 expression in these murine cells is confirmed by MPZL1 knockdown via shRNA and subsequent qPCR mRNA expression analysis (data not shown). **Right panel**: histograms depict FACS data showing the specificity of MPZL1-ChAb by comparing its binding to KrasG12D;p53-/-EV and hMPZL1 OE cell lines. FACS staining is done as described in Figure 3B, using IgG isotype as negative control. Data is acquired utilizing the BD LSRFortessa flow cytometer (BD, Germany) and the BD FACS Diva software v8.0.1m, and analysis is performed with FlowJo v10.

**Suppl. Figure S4.**
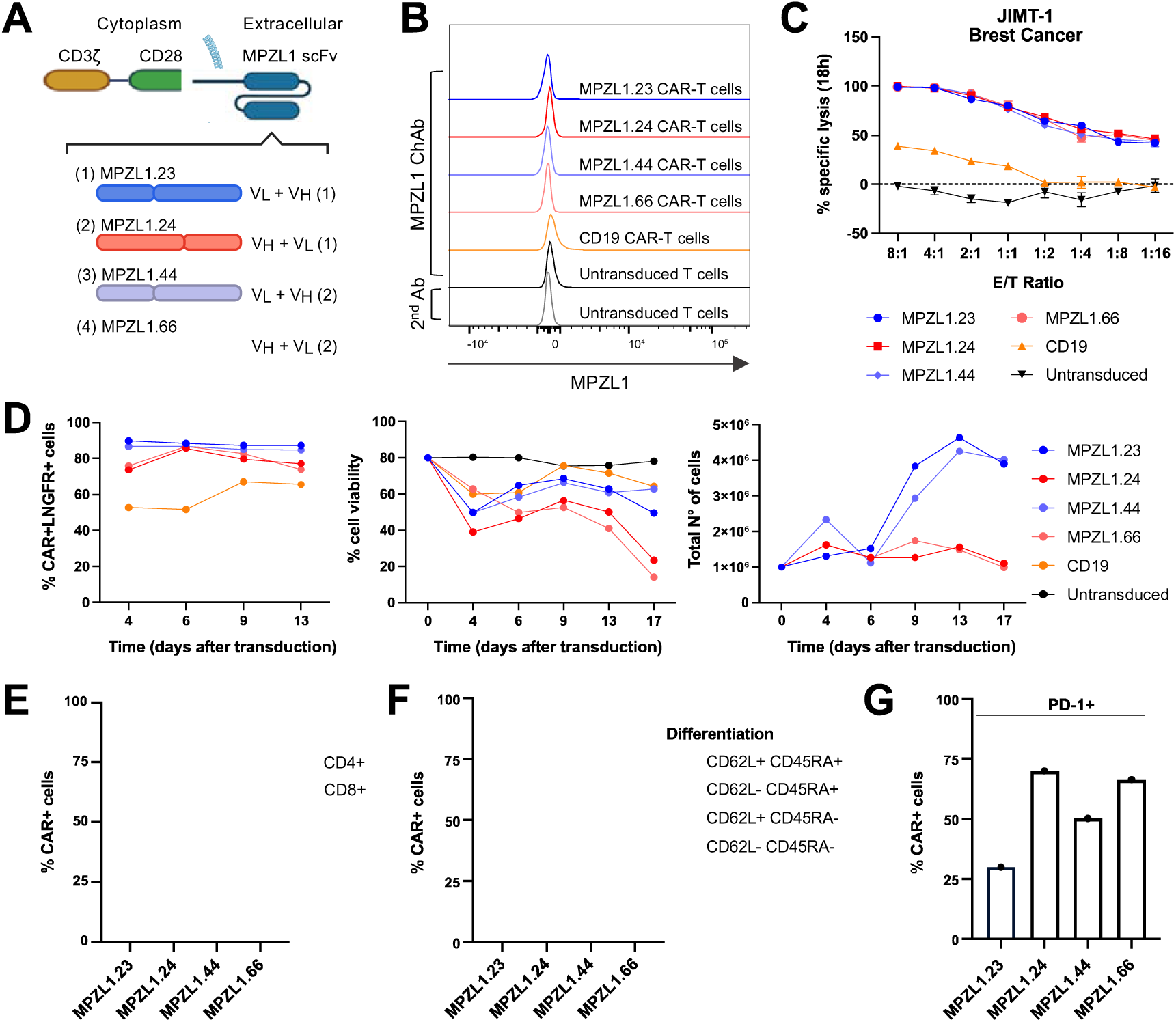
MPZL1 CAR constructs with a V_L_-V_H_ chain configuration persist longer *in vitro*. **(A)** Schematic shows the configuration of the MPZL1 CAR constructs. Created with *BioRender.com*. **(B)** and **(C)** CAR-T cells utilized 6 days after transduction. **(B)** Histograms show FACS data depicting MPZL1-ChAb binding to the different T cell types. The absence of binding excludes CAR-T cell fratricide. Staining is done by incubation with 100 µg/mL of MPZL1-ChAb, followed by incubation with a secondary anti-human antibody coupled a fluorochrome (1:200). **(C)** Bioluminescence-based cytotoxicity assay co-culture during 18 hours of the different CAR-T cells with JIMT-1 breast cancer target cells (MPZL1^high^) at decreasing effector-to-target cell ratios. JIMT-1 cancer cells are stably transduced with a FFLuc-GFP construct. RPMI medium and 0.2% triton X-100 are employed as negative and positive lysis controls, respectively. For the readout, the luciferin substrate is added to the medium, and bioluminescence is detected in a microplate reader. **(D)** CAR-T cell performance over time. **Left panel**: percentage of CAR expression on the cell surface is measured via FACS upon staining with anti-GAM F(ab’)_2_ and LNGFR antibodies. **Middle and right panels**: proliferation is determined by cell counting, and raw numbers (cells/mL) are adjusted to total volumes to express the data in terms of the total N° of cells. Cell viability is determined by exclusion with *Trypan blue*. **(E) –(G)** FACS phenotypic analysis of CAR-T cells 11 days after transduction. **(E)** Percentage of CD4+ and CD8+ CAR-T cells. **(F)** CAR-T cells are classified according to their differentiation pattern determined by CD62L and CD45RA markers. **(G)** Percentage of CAR-T cells expressing the exhaustion marker PD-1. For the antibodies and the staining procedure, check the Methods section. In all cases, acquisition is done using the Cytek^®^ Aurora (Cytek Biosciences, USA) and the SpectroFlow software, while analysis is carried out with FlowJo v10.

**Suppl. Figure S5.**
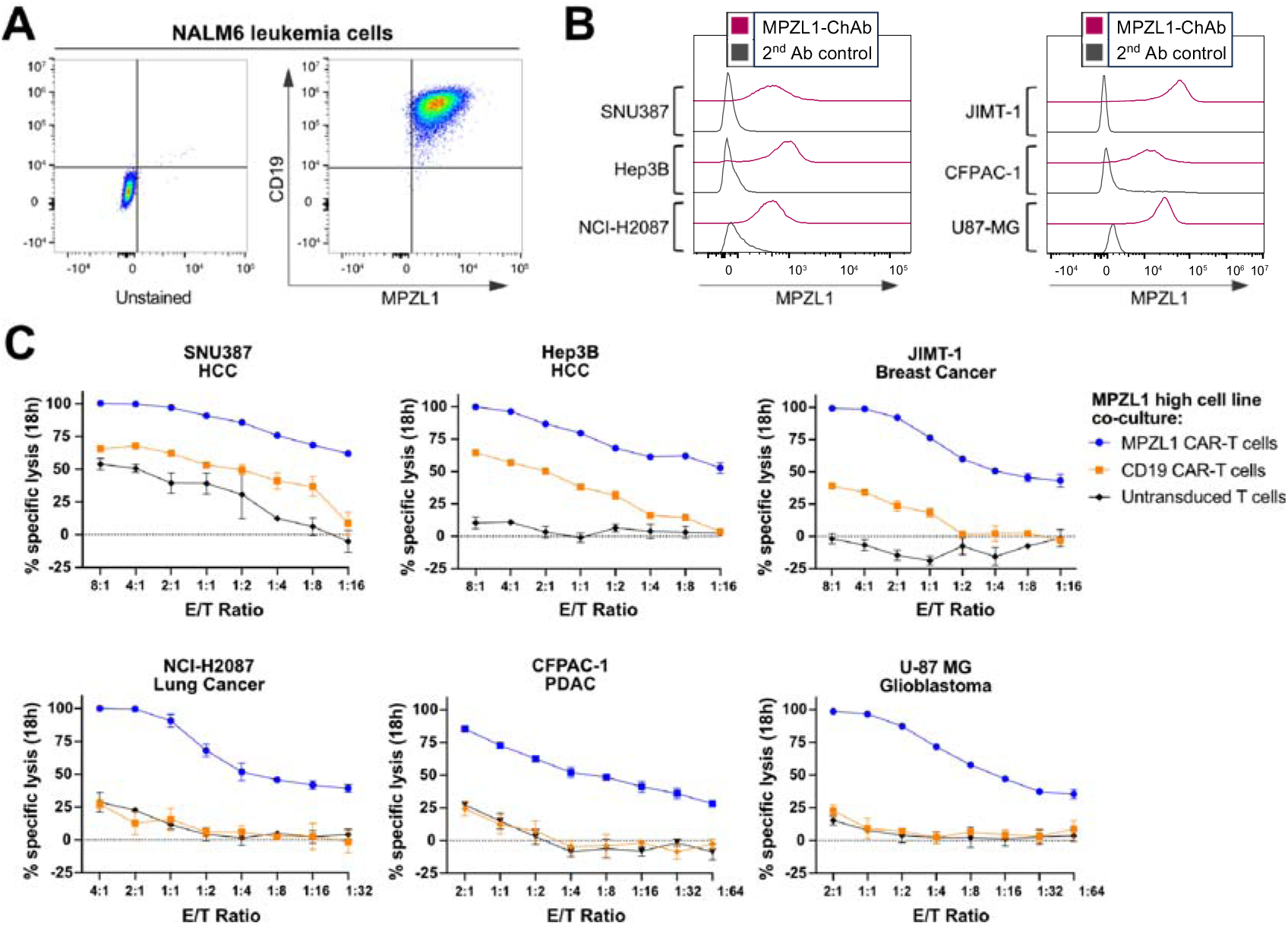
MPZL1 CAR-T cells induce cell lysis in MPZL1^high^ cancer cell lines from various cancer types *in vitro*. **(A)** Dot plots show FACS data depicting NALM6 leukemia cell line (MPZL1^high^; CD19^high^), unstained (left), and double-stained for MPZL1 and CD19. Cells are first stained with MPZL1-ChAb (100 µg/mL), followed by incubation with an anti-human antibody conjugated to a fluorochrome (1:200), and a third staining step with an anti-human PE-CD19 antibody. **(B)** Histograms depict FACS data showing binding of MPZL1-ChAb to cell lines from various tumor entities. Cells are detached with *Versene*, and staining is performed by a first incubation with 100 µg/mL of MPZL1-ChAb, followed by a second incubation with a fluorochrome-coupled secondary anti-human antibody (1:200). The negative control is provided by only using the secondary antibody. In (A) and (B), acquisition is performed using the Cytek^®^ Aurora (Cytek Biosciences, USA) and the SpectroFlow software, while analysis is conducted with FlowJo v10. **(C)** Bioluminescence-based cytotoxicity assay co-culture for 18 hours of MPZL1 CAR, CD19 CAR and untransduced T cells with the MPZL1^high^ cancer cell lines shown in (B). All cancer cells are stably transduced with a FFLuc-GFP construct. RPMI medium and 0.2% triton X-100 are used as negative and positive lysis controls, respectively. For the experimental readout, luciferin substrate is added to the medium, and bioluminescence is detected in a microplate reader.

**Suppl. Figure S6.**
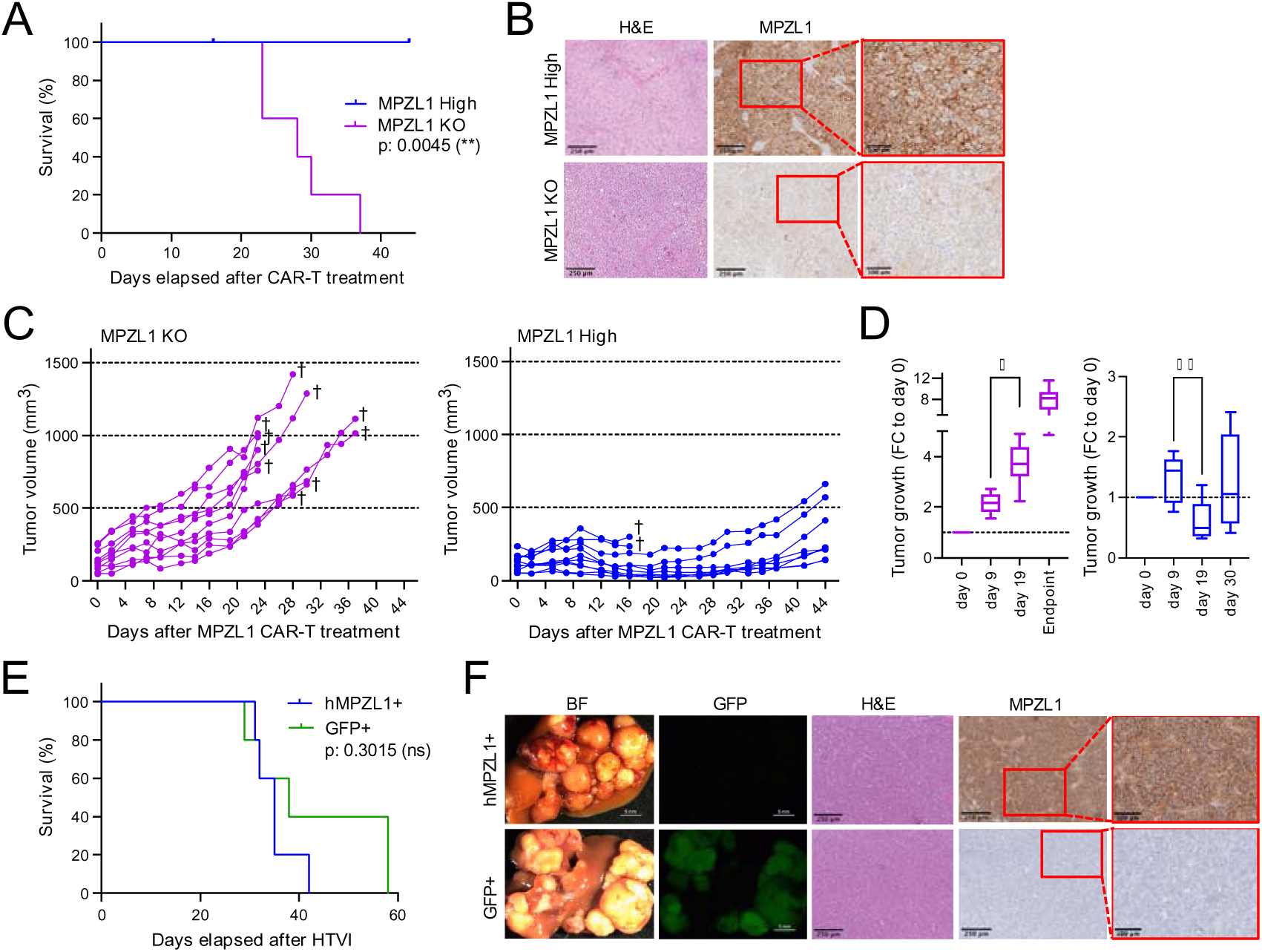
Further in vivo studies. **(A)** – **(D) MPZL1 CAR-T cells exhibit potent effects in human breast cancer xenograft tumors.** Tumor formation in NSG mice is induced by subcutaneous injection of either JIMT-1 EV (MPZL1^high^) or JIMT-1 *MPZL1* KO cells (5×10^6^ cells/flank). A single dose of 10×10^6^ MPZL1 CAR-T cells is applied intravenously when tumors reach an average volume of 120 mm^3^ (n=5 mice/group). **(A)** Kaplan-Meier curve shows survival analysis of the response to MPZL1 CAR-T treatment, with statistics derived from Log-rank test. **(B)** Representative images of FFPE tissue slides from both animal groups: H&E staining (scale bar: 250_μ_m) and MPZL1 IHC staining (two magnifications, scale bars: 250_μ_m and 100_μ_m). **(C)** Tumor growth curves display the volume progression (mm^3^) of *MPZL1* KO tumors (left panel) and MPZL1^high^ tumors (right panel) after therapy administration (day 0). **(D)** Quantification of the tumor growth curves in (C). Boxplots illustrate the size variation of *MPZL1* KO tumors (left panel) and MPZL1^high^ tumors (right panel) at the selected time points relative to treatment administration (day 0) in terms of fold-change (FC), with values >1 representing an increase in tumor volume, and values <1 indicate a decrease thereof. *Endpoint* refers to the volume of each tumor at the time when the respective animal was sacrificed. Statistical analysis performed via one-way ANOVA, and Tukey’s multiple comparisons test (*:p<0.05, **:p<0.01). **(E) – (F) Tumor formation in the HTVI-mediated autochthonous HCC mouse model**. HTVI-mediated tumor induction is performed using the same plasmid cocktail as shown in figure 5F (n=5 mice/group). No treatment applied. **(E)** Kaplan-Meier curve shows survival analysis for both groups of mice in the absence of treatment, with statistics derived from the Log-rank test. **(F)** Characterization of hMPZL1+ and GFP+ tumors. Representative bright-field (BF) and fluorescent GFP stereomicroscopic images of livers (scale bar: 5mm). Representative images of FFPE tissue slides: H&E staining (scale bar: 250_μ_m) and MPZL1 IHC staining (two magnifications, scale bars: 250_μ_m and 100_μ_m).

## Supplementary tables

**Suppl. Table S1.** CSP-coding genes amplified in more than 30% of pan-cancer patient samples (N=90 hits; n=2703 samples).

| CSP gene | Samples amplified (%) | Chromosome | CSP gene | Samples amplified (%) | Chromosome |
| --- | --- | --- | --- | --- | --- |
| *EFNA1* | 33,67 | 1 | *FCRL2* | 32,37 | 1 |
| *EFNA4* | 33,63 | 1 | *CD1B* | 32,33 | 1 |
| *ADAM15* | 33,59 | 1 | *SV2A* | 32,33 | 1 |
| *GBA1* | 33,56 | 1 | *PRRC2C* | 32,33 | 1 |
| *THBS3* | 33,56 | 1 | *ATP1B1* | 32,26 | 1 |
| *ENTREP3* | 33,56 | 1 | *FCRL5* | 32,22 | 1 |
| *PBXIP1* | 33,52 | 1 | *CD1D* | 32,19 | 1 |
| *ADAMTSL4* | 33,33 | 1 | *SELL* | 32,19 | 1 |
| *MPZ* | 33,33 | 1 | *SELP* | 32,19 | 1 |
| *B4GALT3* | 33,30 | 1 | *F5* | 32,19 | 1 |
| *FCGR2A* | 33,22 | 1 | *SLC19A2* | 32,19 | 1 |
| *F11R* | 33,19 | 1 | *SUCO* | 32,11 | 1 |
| *FCGR3A* | 33,07 | 1 | *TNFSF18* | 32,08 | 1 |
| *FCGR3B* | 33,04 | 1 | *KIRREL1* | 32,04 | 1 |
| *LY9* | 33,04 | 1 | *FCGR1A* | 31,85 | 1 |
| *CD244* | 33,04 | 1 | *TOR1AIP2* | 31,52 | 1 |
| *SLAMF7* | 33,04 | 1 | *TOR1AIP1* | 31,52 | 1 |
| *NPR1* | 33,04 | 1 | *QSOX1* | 31,45 | 1 |
| *NCSTN* | 33,04 | 1 | *TOR3A* | 31,26 | 1 |
| *CTSK* | 33,04 | 1 | *MR1* | 31,22 | 1 |
| *SLAMF1* | 33,00 | 1 | *FAM20B* | 31,22 | 1 |
| *IGSF8* | 33,00 | 1 | *ANGPTL1* | 31,15 | 1 |
| *CD48* | 33,00 | 1 | *LAMC1* | 31,08 | 1 |
| *CD84* | 33,00 | 1 | *COLGALT2* | 31,08 | 1 |
| *OLFML2B* | 33,00 | 1 | *LAMC2* | 31,04 | 1 |
| *CGN* | 33,00 | 1 | *EDEM3* | 30,93 | 1 |
| *SLAMF6* | 33,00 | 1 | *HNRNPU* | 30,67 | 1 |
| *FCGR2B* | 33,00 | 1 | *NFASC* | 30,56 | 1 |
| *SSR2* | 33,00 | 1 | *PTGS2* | 30,45 | 1 |
| *GLMP* | 32,96 | 1 | *HMCN1* | 30,37 | 1 |
| *DDR2* | 32,93 | 1 | *PRELP* | 30,11 | 1 |
| *SEMA4A* | 32,93 | 1 | *ADORA1* | 30,04 | 1 |
| *LMNA* | 32,93 | 1 | *ST3GAL1* | 35,15 | 8 |
| *TAGLN2* | 32,93 | 1 | *TNFRSF11B* | 35,04 | 8 |
| *CERS2* | 32,93 | 1 | *COL14A1* | 34,85 | 8 |
| *FCRL6* | 32,85 | 1 | *LRP12* | 34,15 | 8 |
| *FLG2* | 32,70 | 1 | *ANGPT1* | 34,07 | 8 |
| *PEAR1* | 32,63 | 1 | *TRHR* | 33,89 | 8 |
| *CREG1* | 32,59 | 1 | *SLC45A4* | 33,89 | 8 |
| *CADM3* | 32,59 | 1 | *LY6K* | 33,81 | 8 |
| *BCAN* | 32,56 | 1 | *PKHD1L1* | 33,70 | 8 |
| *NTRK1* | 32,56 | 1 | *EEF1D* | 33,56 | 8 |
| *CD1C* | 32,48 | 1 | *CPQ* | 33,04 | 8 |
| *MPZL1* | 32,45 | 1 | *DPY19L4* | 32,89 | 8 |
| *FCRL3* | 32,41 | 1 | *CDH17* | 32,45 | 8 |

**Suppl. Table S2.** Chr. 1q-amplified CSP-coding genes in HCC (N=77 hits, n=370 samples).

| **CSP gene** | **Samples amplified (%)** | **CSP gene** | **Samples amplified (%)** | **CSP genes** | **Samples amplified (%)** |
| --- | --- | --- | --- | --- | --- |
| *ADAM15* | 76.49 | *CD244* | 74.32 | *F5* | 73.51 |
| *EFNA1* | 76.22 | *SLAMF7* | 74.32 | *SLC19A2* | 73.51 |
| *EFNA4* | 76.22 | *CREG1* | 74.32 | *ADAMTSL4* | 73.24 |
| *PBXIP1* | 76.22 | *FCRL5* | 74.32 | *FLG2* | 73.24 |
| *GBA1* | 75.95 | *SUCO* | 74.32 | *SELP* | 73.24 |
| *THBS3* | 75.95 | *SLAMF1* | 74.05 | *ADORA1* | 73.24 |
| *ENTREP3* | 75.95 | *CD48* | 74.05 | *CTSK* | 72.97 |
| *NPR1* | 75.14 | *CD84* | 74.05 | *CERS2* | 72.97 |
| *SSR2* | 75.14 | *TAGLN2* | 74.05 | *PRELP* | 72.97 |
| *GLMP* | 75.14 | *FCRL6* | 74.05 | *EDEM3* | 72.43 |
| *PEAR1* | 75.14 | *FCRL3* | 74.05 | *HNRNPU* | 72.43 |
| *NTRK1* | 75.14 | *PRRC2C* | 74.05 | *TOR3A* | 72.16 |
| *MPZ* | 74.86 | *ATP1B1* | 74.05 | *FAM20B* | 72.16 |
| *FCGR2A* | 74.86 | *TNFSF18* | 74.05 | *ANGPTL1* | 72.16 |
| *FCGR3A* | 74.86 | *NCSTN* | 73.78 | *HMCN1* | 72.16 |
| *FCGR3B* | 74.86 | *IGSF8* | 73.78 | *TOR1AIP2* | 71.89 |
| *FCGR2B* | 74.86 | *SLAMF6* | 73.78 | *TOR1AIP1* | 71.89 |
| *SEMA4A* | 74.86 | *CADM3* | 73.78 | *QSOX1* | 71.89 |
| *LMNA* | 74.86 | *CD1C* | 73.78 | *LAMC1* | 71.89 |
| *BCAN* | 74.86 | *FCRL2* | 73.78 | *COLGALT2* | 71.89 |
| *B4GALT3* | 74.59 | *CD1B* | 73.78 | *LAMC2* | 71.89 |
| *OLFML2B* | 74.59 | *CD1D* | 73.78 | *MR1* | 71.62 |
| *DDR2* | 74.59 | *KIRREL1* | 73.78 | *PTGS2* | 71.62 |
| *MPZL1* | 74.59 | *NFASC* | 73.78 | *SV2A* | 68.65 |
| *F11R* | 74.32 | *CGN* | 73.51 | *FCGR1A* | 64.05 |
| *LY9* | 74.32 | *SELL* | 73.51 |  |  |

**Suppl. Table S3.** Chr. 1q-amplified CSP-coding genes differentially expressed between cancer and healthy liver tissues (N=19 hits; n=369 tumor and 160 normal samples). mRNA differential expression is considered as |Log_2_FC|>1.

| Gene | Log<sub>2</sub>FC | Gene | Log<sub>2</sub>FC | Gene | Log<sub>2</sub>FC |
| --- | --- | --- | --- | --- | --- |
| *GBA* | 2.452 | *PBXIP1* | 1.440 | *NCSTN* | 1.196 |
| *LAMC1* | 2.038 | *ATP1B1* | 1.370 | *F5* | 1.152 |
| *LMNA* | 1.913 | *FAM189B* | 1.313 | *FAM20B* | 1.114 |
| *GLMP* | 1.742 | *ADAM15* | 1.286 | *SUCO* | 1.113 |
| *TAGLN2* | 1.716 | *EFNA4* | 1.255 | *IGSF8* | 1.056 |
| *SSR2* | 1.633 | *MPZL1* | 1.247 | *TOR3A* | 1.026 |
| *OLFML2B* | 1.568 |  |  |  |  |

**Suppl. Table S4.** Candidate CSPs for a targeted therapy (N=6 hits).

| Gene | Full name | Protein cellular localization |
| --- | --- | --- |
| *ATP1B1* | Na<sup>+</sup>/K<sup>+</sup> transporting ATPase subunit beta-1 | Cell membrane |
| *FAM189B* | Endosomal transmembrane epsin interactor 3 | Membrane (*ENTREP3*) |
| *EFNA4* | Ephrin-A4 | Cell membrane |
| *MPZL1* | Myelin protein zero-like 1 | Membrane / Cell surface |
| *IGSF8* | Immunoglobulin superfamily member 8 | Cell membrane |
| *ADAM15* | Disintegrin and metalloproteinase domain-containing protein 15 | Plasma membrane / Endomembrane system |

## Notes

### Competing Interest Statement

The authors have declared no competing interest.

